# Local glycan engineering induces systemic antitumor immune reactions via antigen cross-presentation

**DOI:** 10.64898/2026.05.04.720097

**Authors:** Natalia Rodrigues Mantuano, Michael T. Sandholzer, Emiel Rossing, Johan F.A. Pijnenborg, Andreas Zingg, Filip Filipsky, Ronja Wieboldt, Amanda Carlos Paulino, Isabelle V. Menegoy Siqueira, Thomas J. Boltje, Heinz Läubli

## Abstract

Immune checkpoint inhibitors (ICI) have revolutionized cancer therapy, yet response rates remain suboptimal across many solid tumors, and resistance mechanisms, particularly those involving glycans, are not fully understood. Recent studies have identified sialic acid-containing glycans and their interactions with Siglec receptors on tumor-associated macrophages as an important contributor to immune suppression within the tumor microenvironment (TME). Targeting this sialic acid–Siglec axis by glycan engineering with sialidases and other glycosidases has shown therapeutic potential in preclinical models. However, safe and effective delivery of sialidases to tumors remains a challenge. Here, we present a novel approach using adeno-associated virus (AAV)-mediated therapy to deliver sialidases (AAV^Sia^) and other glycosidases, including fucosidase, directly to the TME. Intratumoral administration of AAV^Sia^ in mouse models resulted in significant tumor growth reduction, enhanced survival, and robust systemic antitumor immunity through improved cross-presentation and dendritic cell activation. Furthermore, combining local sialidase expression with fucosidase treatment and classical PD-1 blockade allowed a synergistic effect, amplifying antitumor response. Our findings highlight the therapeutic promise of glycoengineering the TME using local delivery systems and support the development of combination strategies to overcome glycan-mediated resistance in cancer immunotherapy.

**Graphical abstract:** 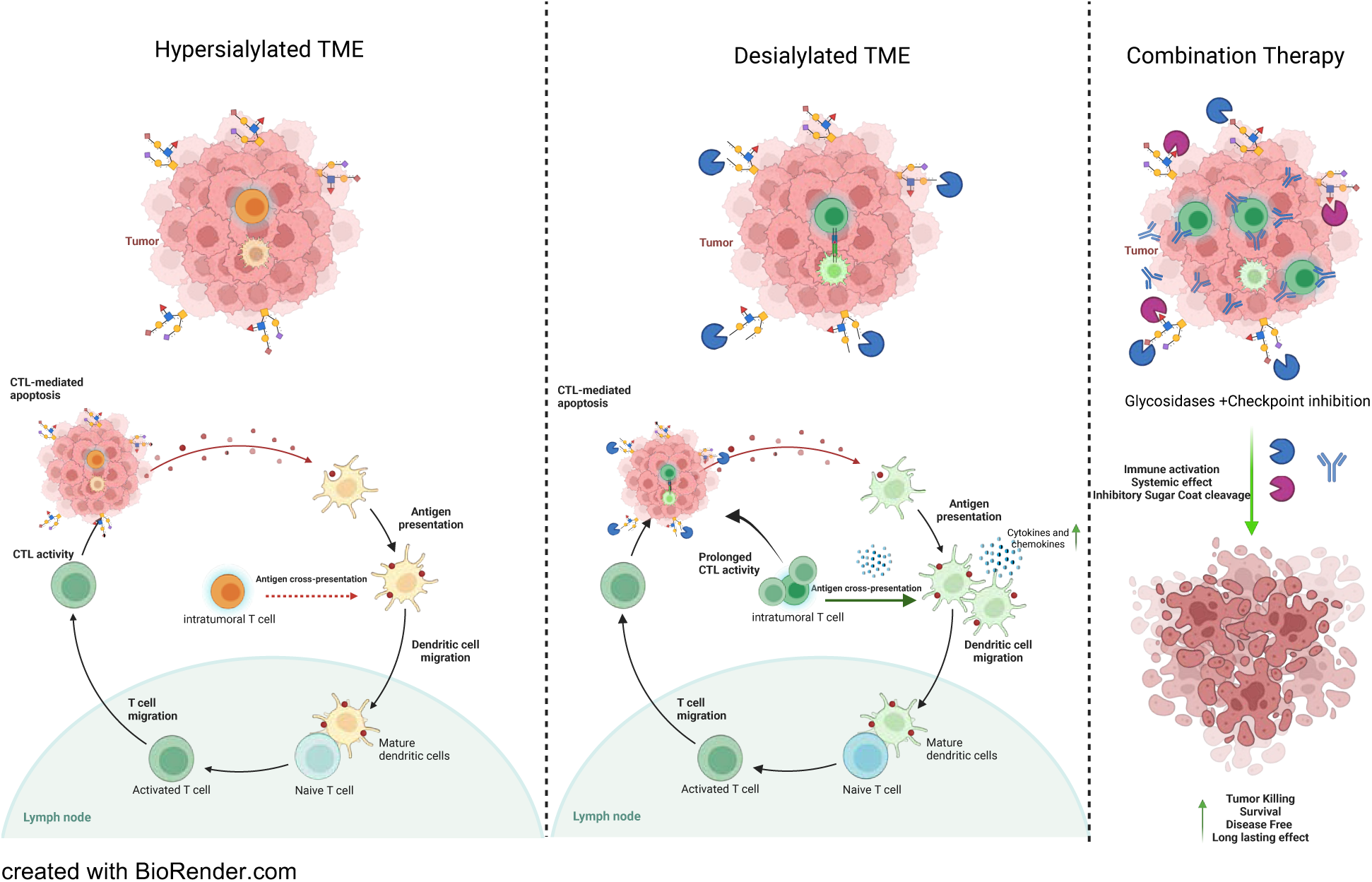

## Introduction

The advent of immune checkpoint inhibition (ICI) has brought about a transformative shift in the available treatment approaches for cancer patients. Nevertheless, despite these advances, the overall response rates across most solid tumor types remain modest, typically ranging between 10% and 50%. Moreover, the mechanisms underlying resistance to ICI are still not fully understood^1, 2^. Consequently, further studies are required to elucidate these resistance pathways. Recent evidence suggests that alterations in glycosylation may contribute to treatment resistance, highlighting the need to develop strategies to overcome glycan-associated resistance^3, 4^.

In recent years, significant progress has been made in understanding the role of sialic acid-containing glycans and their interaction with sialic acid-binding immunoglobulin-like lectin (Siglec) receptors in the tumor microenvironment ^5, 6, 7^. These interactions have emerged as a promising area of research, potentially serving as a novel immune checkpoint and a viable target for cancer immunotherapy. Myeloid cells, including tumor-associated macrophages (TAMs) and dendritic cells, have been identified as the main mediators of sialic acid-mediated immune suppression through Siglec receptors ^8^. The sialic acid-Siglec pathway can be disrupted by different approaches, including the use of blocking antibodies, sialidases, and glycomimetics ^9, 10, 11^. Sialidases that are directed to the tumor microenvironment have been shown to significantly reduce tumor growth in different pre-clinical models ^6, 8^ ^12^, and a first clinical trial using a bi-sialidase fusion protein was completed in 2024 (GLIMMER-01 Trial, NCT05259696).

However, a critical concern remains whether sufficiently high intra-tumoral sialidase levels can be attained in patients without inducing systemic toxicity. To address this challenge, viral gene therapy employing recombinant adeno-associated viruses (AAV) could be used to increase local concentrations of effectors. AAV therapy possesses several advantageous features, including low immunogenicity, minimal side effects, sustained transgene expression, adaptability across various cancer types, and no direct association with human disease pathogenesis ^13^. These attributes position AAV therapy as a compelling candidate for targeted and safe delivery of therapeutic agents in cancer treatment strategies ^14, 15, 16^.

In this study, we engineered an AAV encoding a H1N1 Influenza A sialidase (AAV^Sia^), and evaluated its therapeutic potential in mouse models. Our results demonstrate that intratumoral administration of AAV^Sia^ significantly improves survival rates and elicits robust antitumor immunity, primarily mediated through cross-presentation mechanisms, increasing dendritic cell migration and activation. Importantly, we observed systemic immune responses enhanced specific anti-tumor T cell activity following local AAV^sia^ treatment. Finally, we demonstrate that local sialidase expression can be combined with other glyco-engineering strategies, including fucosidase treatment within the TME, and enhance the efficacy of PD-1 blockade. This combined approach can synergistically target the tumor microenvironment, amplifying antitumor immune responses. These findings highlight the promising immunotherapeutic potential of tumor microenvironment glycoengineering in cancer treatment.

## Results

### Engineering an AAV-based delivery platform for glycosidases

We designed a recombinant AAV type-2 virus encoding a H1N1 influenza A sialidase (AAV^Sia^), under the control of a cytomegalovirus (CMV) promoter (**Fig.1a**). Influenza A sialidases/neuraminidases are naturally expressed intravirion, containing a transmembrane domain that targets the enzyme to the endoplasmic reticulum and facilitates its membrane integration and transport to the plasma membrane, where the head domain then cleaves off sialic acid, promoting virus release ^17, 18^. Single intra-tumoral (i.t.) treatment with AAV^Sia^ compared with intravenous (i.v.) shows that local i.t. injection in MC38 solid tumors has higher efficacy (**Fig.1b,c and Supplementary Fig.1a**). Compared with intratumoral (i.t.) injection, systemic intravenous (i.v.) administration results in less efficient tumor delivery (**Supplementary Fig. 1b**). Local i.t. administration allow higher amounts of AAV to reach the tumor while minimizing exposure to off-target organs such as the brain and liver (**Fig. 1d and Supplementary Fig. 1b**). Consistently, transmembrane expression was detected within the tumor microenvironment (**Fig. 1e,f**), whereas sialidase remained undetectable in the plasma of locally treated mice (**Supplementary Fig. 1c,d**), thereby mitigating systemic toxicity and desialylation of distant tissues. Treatment with AAV^Sia^ intratumoral results in sialidase expression (**Fig. 1e,f**) and desialylation (**Fig. 1g, h**) both 1 and 2 weeks after a single injection, as evidenced by SNA (*Sambucus Nigra*) lectin staining. Overall, the findings suggest that an AAV-based delivery platform for expressing glycan-cleaving glycosidases such as sialidase holds promise for cancer therapy, offering effective local treatment with minimal off-site effects.

**Figure 1.**
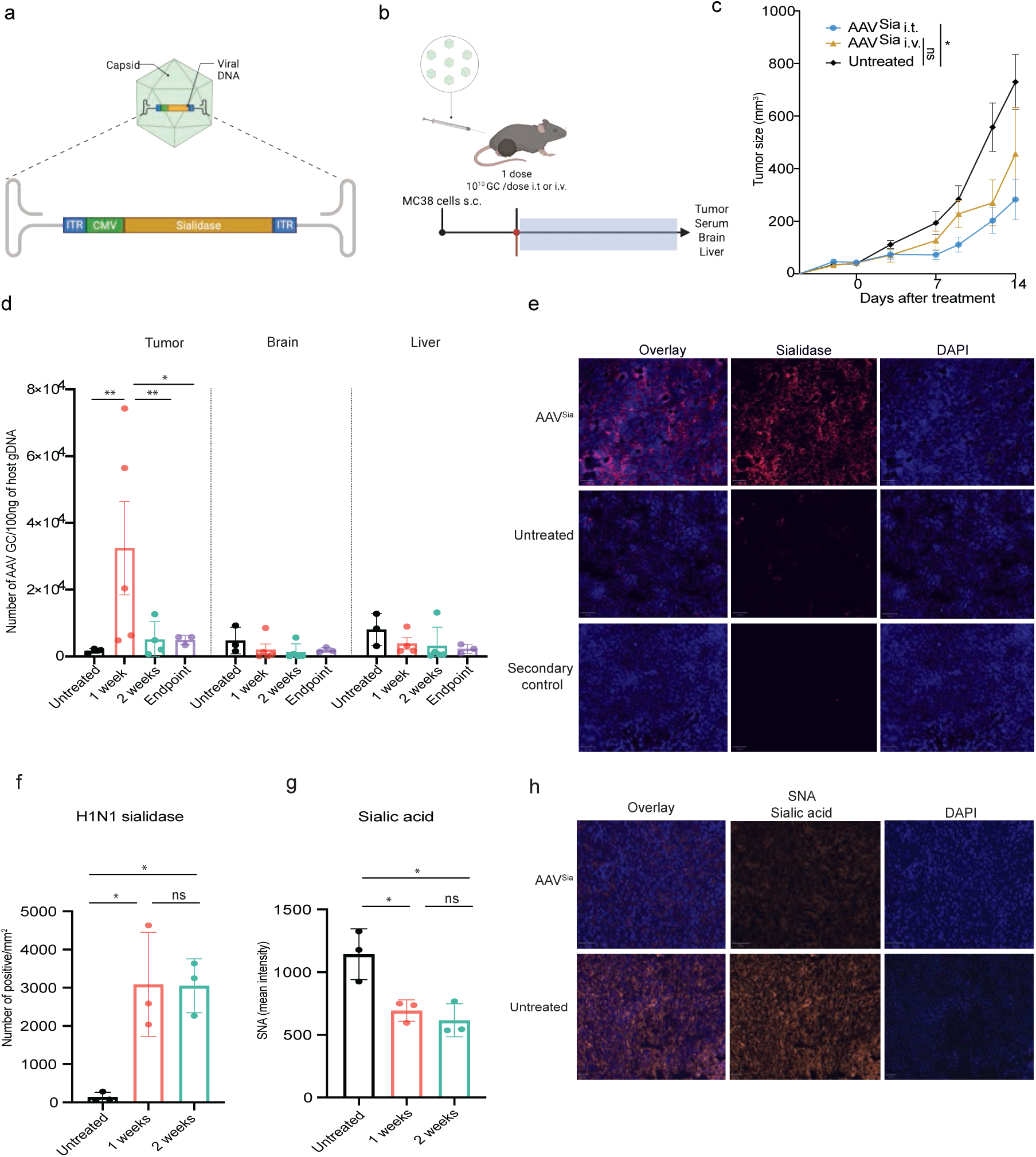
– Virus-based delivery platform for sialidase expression. **a,** Schematic of recombinant adeno-associated virus (AAV) type 2 engineered to express Influenza A H1N1 sialidase/neuraminidase (AAVSia). **b,** MC38 tumors and treatment scheme. MC38 cells injected subcutaneously (s.c.) and treated with AAV^Sia^ intratumorally (i.t.) or intravenously (i.v.) once tumors reached a defined size. Tumors, serum, brain, and liver harvested at one week, two weeks, or experimental endpoint. **c,** Tumor growth curves measured three times per week; tumor volume (mm³) calculated for each treatment group (n=5-8). **d,** AAV detection by quantitative PCR (qPCR) in tumor, brain, and liver at one week, two weeks, or endpoint (n=3-4). **e,f,** Immunofluorescence detection of H1N1 sialidase in i.t.-treated tumors (n=3). **g,h,** Sialic acid levels assessed by SNA (*Sambucus nigra*) staining in i.t.-treated tumors (n=3). Significance determined by two-way ANOVA (**c,d**) or one-way ANOVA (**f,g**), and Tukey’s multiple comparison test. *P < 0.05, **P < 0.01. **a,b** were Created with BioRender.com.

### Local sialidase expression suppresses tumor growth and shapes the tumor immune microenvironment

Sialidase treatment has been previously recognized for its efficacy against cancer and its ability to stimulate the immune system by evading engagement with inhibitory receptors like Siglecs^6, 19, 20^. To further explore the therapeutic potential of AAV^Sia^ in various solid tumor models amenable to local treatment, we conducted a series of experiments. As controls, we generated a loss-of-function enzyme, AAV^LOF^, and a virus without any payload (AAV^null^). AAV^Sia^ administration was initiated when tumors reached ∼50 mm³ and delivered intratumorally at 10¹⁰ genomic copies (GC) per injection every 2–3 days for four doses. Control treatments were administered under the same conditions (**Fig. 2a**)

**Figure 2.**
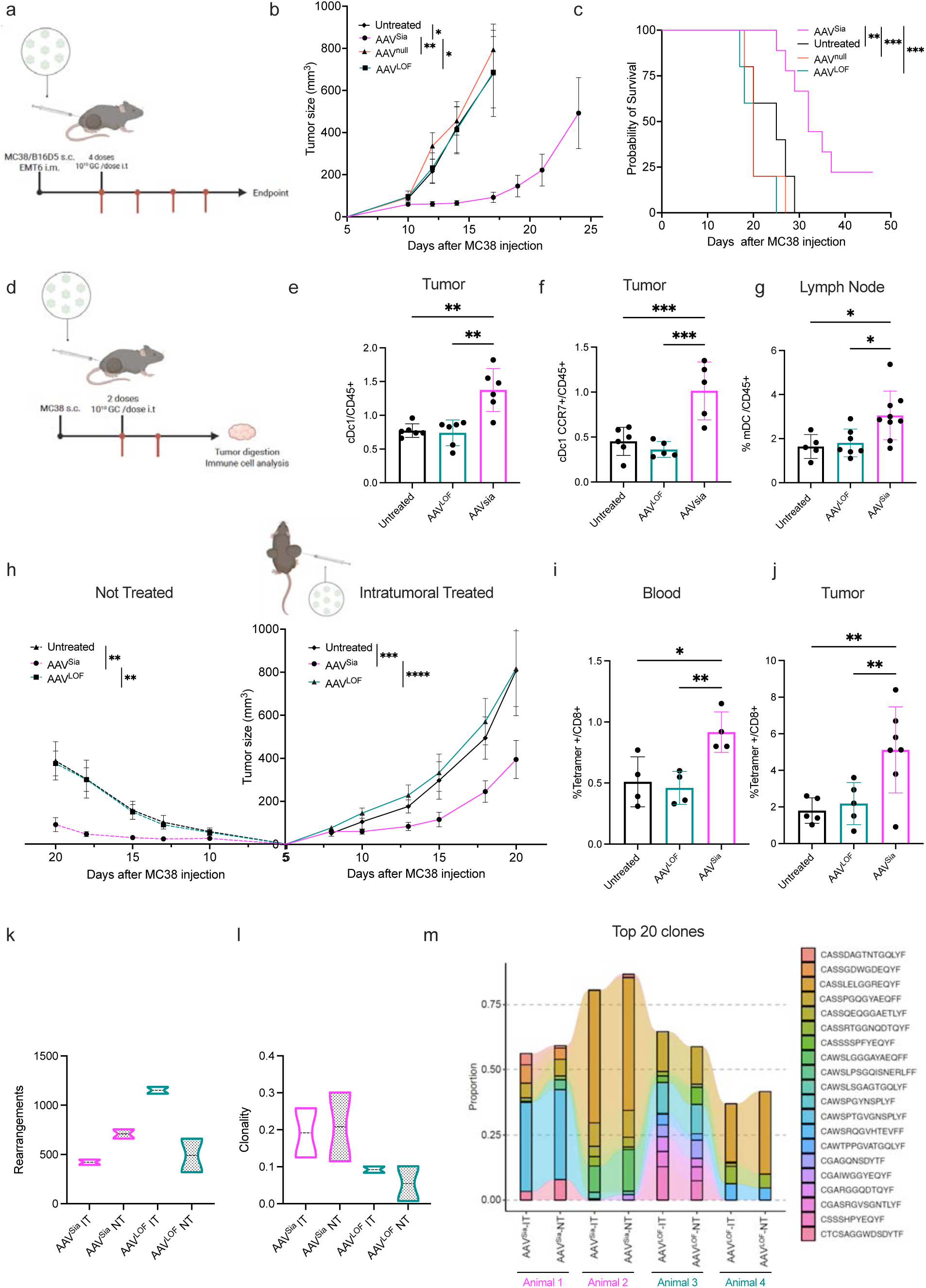
– Local Sialidase Expression Suppresses Tumor Growth and Enhances Systemic Antitumor Activity. **a,** Treatment scheme. AAV^Sia^ administration initiated when tumors reached ∼50 mm³. Virus diluted in PBS and delivered intratumorally (i.t.) at 10¹⁰ genomic copies (GC) per injection every 2–3 days for four doses. AAV^LOF^ or AAV^null^ administered under the same conditions as controls. **b,** Tumor volume measured every 2–3 days with calipers until the humane endpoint.**c,** Survival relative to the tumor volume endpoint. **d,** MC38 tumors and lymph nodes collected one week after two treatment doses for immune infiltrate analysis. **e,f,** Frequencies of conventional type 1 dendritic cells (cDC1s) and CCR7⁺ cDC1s. **g,** Migratory dendritic cells (migDCs) in lymph nodes. **h,** Abscopal tumor model. MC38 cells implanted in the right flank and, three days later, in the left flank. Bilateral tumor growth with tumors intratumorally treated (IT) with AAVSia or AAV^LOF^ or left untreated (NT) in the same mice. Untreated control mice (both tumors NT) were included. **i,j,** Tetramer staining for the Adpgk peptide (MC38 neoantigen) in CD8⁺ T cells in blood and tumors analyzed by flow cytometry. **k–m,** TCRβ sequencing of IT and NT tumors from mice treated with AAV^Sia^ or AAV^LOF^. Growth curves are shown as mean ± S.E.M. (n = 6–11). Bar plots are presented as mean ± S.D. (n = 4–6). Violin plots show the distribution with the mean indicated by a dotted line (n = 2). Statistical significance among multiple groups was assessed using one-way ANOVA (bar plots) or two-way ANOVA (growth curves), and Tukey’s multiple comparison test. Survival curves were analyzed using the Kaplan–Meier method (e). *P < 0.05, **P < 0.01, ***P < 0.001. ****p< 0.0001. **a,d** were created with BioRender.com.

In subcutaneous models utilizing MC38 and B16D5 cells, we observed a decrease in tumor growth and a significant increase in overall survival upon treatment with AAV^Sia^ compared to control groups (**Fig. 2b,c and Supplementary Fig. 2a,b**). In the MC38 tumor model, a fraction of AAV^Sia^-treated mice remained tumor-free after treatment. Upon re-challenge with tumor cells in the left flank, no tumor relapse was observed compared to newly injected control mice (**Supplementary Fig. 2c**). Similar results were obtained in the orthotopic EMT6 breast cancer model, in which treatment with AAV^Sia^ decreased tumor growth and enhanced survival (**Supplementary Fig. 2d,e**).

To assess the effect of circulating antibodies against influenza sialidase on the efficacy of AAV^Sia^, we vaccinated mice and subsequently injected them with MC38 cells subcutaneously (**Supplementary Fig. 2f**). Circulating anti-H1N1 sialidase antibodies were confirmed in vaccinated animals (**Supplementary Fig. 2g**). We confirmed there was no difference in tumor growth and survival upon vaccination and subsequent AAV^Sia^ treatment (**Supplementary Fig. 2h,i**).

To investigate changes in immune infiltration following AAV^Sia^-mediated reduction of sialic acid levels, a short treatment regimen consisting of two intratumoral doses of AAV^Sia^ or controls was performed. One week later, tumors and lymph nodes were harvested and processed for immune cell profiling (**Fig. 2a**). Although desialylation was observed in the TME (**Fig. 1 g,h**), sialidase expression was detected on the surface of CD45⁻ intratumoral cells, but not in CD45⁺ immune infiltrated, following AAV^Sia^ and AAV^LOF^ treatment (**Supplementary Fig. 2j**).

AAV^Sia^ treatment increased the infiltration of conventional type 1 dendritic cells (cDC1s), including the CCR7⁺ cDC1 subset (**Fig. 2e,f**), whereas no differences were observed in conventional type 2 dendritic cells (cDC2s) (**Supplementary Fig. 2k**). In draining lymph nodes, the frequency of migratory dendritic cells (migDCs) was increased, while resident dendritic cells were decreased compared to untreated mice, but not relative to AAV^LOF^-treated controls (**Fig. 2g and Supplementary Fig. 2n**). cDC1s from AAV^Sia^-treated tumors exhibited increased expression of the activation marker CD80 and higher intracellular IL-12 levels (**Supplementary Fig. 2l,m**).

CD8⁺ T cells were increased and displayed enhanced proliferation, as indicated by Ki67 expression, compared to control groups (**Supplementary Fig. 2o,p**). In contrast, total CD3⁺ and CD4⁺ T cell infiltration remained unchanged (**Supplementary Fig. 2q,r**). Macrophage infiltration was also unchanged, however, MHC II expression, indicative of an antitumoral M1-like polarization state, was increased following AAV^Sia^ treatment compared with AAV^LOF^ (**Supplementary Fig. 2s,t**).

Taken together, these results suggest that immune mechanisms, including enhanced T cell responses and increased cDC1 activation and migration mediate the anti-tumor effects of AAV^Sia^ treatment. Furthermore, the absence of tumor relapse upon re-challenge demonstrates the induction of an anti-tumor memory response post-treatment.

### Intratumoral AAV^Sia^ promotes systemic antitumor activity through desialylation

Next, we wanted to verify whether local desialylation induces systemic antitumoral effects and identify underlying cellular and molecular mechanisms. A screening of 44 plasma cytokines and chemokines revealed that AAV^Sia^ treatment decreased pro-tumoral factors, including TNF-α, IL-10, M-CSF, and CCL5, compared with untreated controls, whereas only IL-10 and M-CSF were significantly reduced relative to AAV^LOF^ (**Supplementary Fig. 3b**). Notably, tumor-suppressive factors involved in T cell recruitment, including IL-16, CX3CL1, and CXCL9, were increased following AAV^Sia^ treatment, while CXCL10 was elevated only in comparison with untreated mice (**Supplementary Fig. 3b**).

To access both local and distant effects triggered by local desialylation following AAV^Sia^ treatment, we employed the MC38 tumor model to study potential abscopal effects. In this experimental setup, cancer cells were injected into the right flank (intratumoral treatment, IT), followed by injection into the left flank (not treated, NT) after 3-4 days. tumoral injections (IT), and the tumors on the left flank remained untreated (NT). IT tumors treated with AAV^Sia^ showed reduced growth compared with controls (**Fig. 2h**). The NT tumors were also affected by treatment of the primary tumor (**Fig. 2h**), suggesting a systemic immune response. To exclude direct effects of the virus on the contralateral NT tumor, sialylation was measured using SNA staining. Contralateral virus injection did not induce desialylation in NT tumors compared to IT tumors (**Supplementary Fig. 3c,d**).

Next, MC38-specific tetramer staining using the Adpgk peptide, a MC38 neoantigen ^21^, was performed in blood and contralateral NT tumors to quantify tumor-specific CD8⁺ T cells (**Fig. 2i,j**). Tetramer analysis revealed an increase in circulating tumor-specific CD8⁺ T cells following AAV^Sia^ intra-tumoral treatment compared to untreated and AAV^LOF^ controls. Indeed, intra-tumoral tetramer⁺ CD8⁺ T cells were also elevated in the AAV^Sia^ tumors (**Fig. 2i,j**)

TCR sequencing of T cell clones from IT and NT tumors was performed. AAV^Sia^-treated mice exhibited reduced TCR diversity (**Fig. 2k**) and increased clonal expansion compared with AAV^LOF^-treated controls (**Fig. 2l**). Consistently, higher clonality was observed in both IT and NT tumors from AAV^Sia^-treated mice (**Fig. 2l**). Analysis of the top clones further revealed expanded T cell clones in AAV^Sia^-treated mice (**Fig. 2m)** Additionally, a higher number of shared clonotypes was observed between IT and NT tumors in AAV^Sia^-treated mice. In contrast, AAV^LOF^-treated mice exhibited fewer shared clonotypes between IT and NT tumors (**Supplementary Fig. 3e**)

These findings indicate that local tumor desialylation leads to a systemic immune response, potentially affecting sites not treated intratumorally through increased tumor-specific responses and clonal expansion.

### Virus-based desialylation improves cancer immunotherapy by antigen cross-presentation

To further dissect immune mechanisms involved in effects observed, we employed genetically modified mouse models. Batf3^-/-^ mice lack cDc1 and are defective in cross-presentation ^22^. Tumor-bearing Batf3^-/-^ mice were treated with AAV^Sia^ and compared to wild-type ^4^ mice control (**Fig. 3a**). The results revealed a loss of treatment AAV^Sia^ efficacy in Batf3^-/-^ mice (**Fig. 3b**), indicating that cross-presentation might be important for the efficacy of sialidase treatment. CD11c-DTR mice ^23^ allow for CD11c^+^ cells transient depletion upon diphtheria toxin (DT) injection (**Fig. 3a,c**). Similar findings as in Batf3^-/-^mice were observed in CD11c-DTR mice, where AAV^Sia^ treatment responsiveness was lost when mice were depleted of dendritic cells (**Fig. 3c**). To test the impact of sialic acid loss during cross-presentation, we utilized XCR1^+^ bone marrow-derived dendritic cells (BMDCs), pulsed with irradiated MC38-OVA B2m^-/-^ cells and co-cultured with OT-I cells. Each cell type (dendritic cells or T cells) was pre-treated with soluble H1N1 Influenza A sialidase (Sia) before co-culture to assess the effects of sialic acid loss during cross-presentation (**Fig. 3d**). Lower sialic acid levels in T cells did not significantly affect T cell proliferation during cross-presentation (**Fig. 3e**). The most pronounced effect on cross-presentation was observed when XCR1^+^ BMDCs were desialylated (**Fig. 3e**). As a control, heat-inactivated (HI) enzyme was used, with no changes in T cell proliferation (**Supplementary Fig. 4a**).

**Figure 3.**
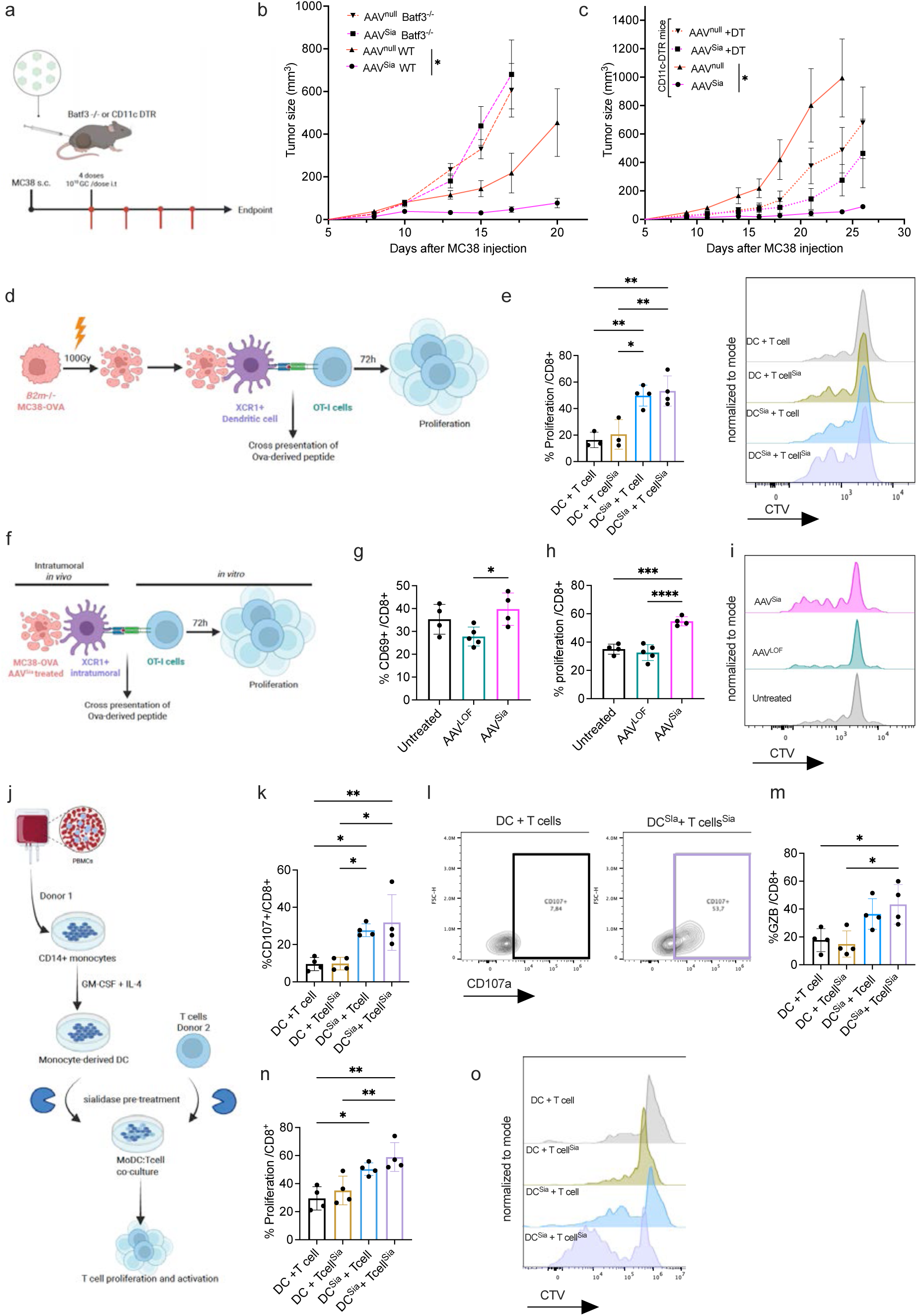
– Virus-based desialylation improves cancer immunotherapy by antigen cross-presentation.**a,** Treatment scheme. MC38 tumor–bearing Batf3⁻/⁻ or CD11c-DTR mice were treated with four intratumoral injections of AAV^Sia^, AAV^null^, or left untreated. **b,** Tumor growth curves of MC38-bearing Batf3⁻/⁻ mice (dashed lines) compared with wild-type (WT; solid lines) controls following treatment. **c,** Tumor growth curves of CD11c-DTR MC38-bearing mice treated with AAV, with (dotted line) or without diphtheria toxin (DT; solid lines).**d,** *In vitro* cross presentation assay using irradiated B2m⁻/⁻ MC38-OVA cells and bone marrow–derived dendritic cells (BMDCs), sorted for XCR1⁺ dendritic cells, and co-incubated with OT-I CD8⁺ T cells for 72 hours following pre-treatment with soluble H1N1 sialidase (Sia).**e,** Quantification of T cell proliferation in the *in vitro* cross-presentation assay, with representative histogram plots.**f,** *In vitro* cross-presentation assay. MC38-OVA–bearing mice were treated intratumorally with AAV^Sia^, AAV^LOF^, or left untreated. One week after two treatment doses, XCR1⁺ intratumoral cells were enriched and co-incubated with OT-I CD8⁺ T cells. **g,** Frequency of CD69⁺ cells among CD8⁺ T cells.**h,** Proliferation of OT-I CD8⁺ T cells in the cross-presentation assay.**i,** Representative histogram plots of Cell Trace Violet (CTV) dilution.**j,** Mixed leukocyte reaction (MLR) scheme. Monocyte-derived dendritic cells (moDCs) were pre-treated with or without sialidase and co-cultured for 5 days with allogeneic CD8⁺ T cells, which were also pre-treated or not with sialidase, at a 1:1 ratio. **k,** Percentage of CD107a⁺ CD8⁺ T cells. **l,** Representative contour plots of CD107a expression. **m,** Percentage of granzyme B (GZB)–positive CD8⁺ T cells. **n,** Percentage of proliferating CD8⁺ T cells.**o,** Representative dot plots of CTV dilution.Growth curves are shown as mean ± S.E.M. (n = 4-11). Bar plots are presented as mean ± S.D. (n = 3–4). Statistical significance among multiple groups was assessed using one-way ANOVA (bar plots) or two-way ANOVA (growth curves), and Tukey’s multiple comparison test. *P < 0.05, **P < 0.01, ***P < 0.001. **a,d,f,g** were created with BioRender.com.

To corroborate the enhanced cross-presentation observed *in vitro* upon AAV^Sia^ treatment, we treated MC38-OVA bearing mice, sorted XCR1^+^ dendritic cells (cDC1), and co-cultured them with OT-I cells *in vitro* (**Fig. 3f**). This resulted in augmented proliferation and CD69 positivity of OT-I cells upon AAV^Sia^ treatment. (**Fig. 3g,h,i**). No changes were observed in CD25^+^ OT-I cells among the treatment groups (**Supplementary Fig. 4b**).

Next, human monocyte-derived dendritic cells (moDcs) were utilized to conduct a mixed lymphocyte reaction, using T cells from a different donor (**Fig. 3j**). Both cell types were isolated from healthy donor PBMCs (peripheral blood mononuclear cells). T cells, moDcs, or both were pre-treated with Sia, and after co-incubation, T cell proliferation and activation were assessed. Increased degranulation (CD107a^+^ cells) and granzyme B+ cells were observed only when moDCs were treated with Sia or both cell types were desialylated (**Fig. 3k,l,m**). Intracellular interferon-gamma and PD-1 levels on the surface of T cells did not change among the treatment groups (**Supplementary Fig. 4c,d)**. Our results indicate that desialylation of moDCs is crucial for enhanced T cell proliferation (**Fig. 3n,o**). Sialidase pretreated moDCs were co-incubated with or without LPS, followed by analysis of maturation markers. CD80 expression was enhanced upon sialidase treatment during LPS-induced maturation, whereas CD86 was increased only in immature moDCs pre-treated with sialidase (**Supplementary Fig. 4e,f**). CD40 expression remained unchanged (**Supplementary Fig. 4g**).

Together, our results suggest that enhanced cross-presentation occurs upon local desialylation mediated by cDC1 cells. In addition, desialylation of T cells alone does not appear to significantly enhance cross-presentation. Using human moDCs, we found that allogeneic (non-self) responses were amplified when moDCs exhibited reduced levels of sialic acid, as reflected by increased T cell proliferation and degranulation.

### Sc-RNAseq shows enhanced cytokine and migration-associated activation in cDC1s upon treatment with AAV^sia^

To further investigate the role of cDC1 cells, single-cell RNA sequencing (scRNA-seq) was performed on CD11c⁺ F4/80^-^ cells sorted from MC38-OVA tumors treated with AAV^Sia^, AAV^LOF^, or untreated. We identified distinct populations of dendritic cells, macrophages, and monocytes (**Supplementary Fig. 5a,b**). Dendritic cell subclusters were further subdivided into proliferating dendritic cells, cDC2, and migratory DCs (migDCs) (**Fig. 3a and Supplementary Fig. 5c,d**). CITE-seq was performed to assess the expression of the extracellular markers XCR1 (a cDC1 marker) and CCR7 (a migration marker) on protein level, and analyzed alongside RNA expression levels (**Supplementary Fig. 5d**).

Genes related to antigen presentation and costimulation (*Ciita, Psme3, Cd40, Cd86, Cd80,* and *Wdfy4*) were enhanced in the AAV^Sia^ group when analyzing the cDC1 population (**Fig. 4b,c**). On the other hand, transporters associated with the antigen-processing complex, *Tap1* and *Tap2*, were increased in the AAV^LOF^ group.

**Figure 4.**
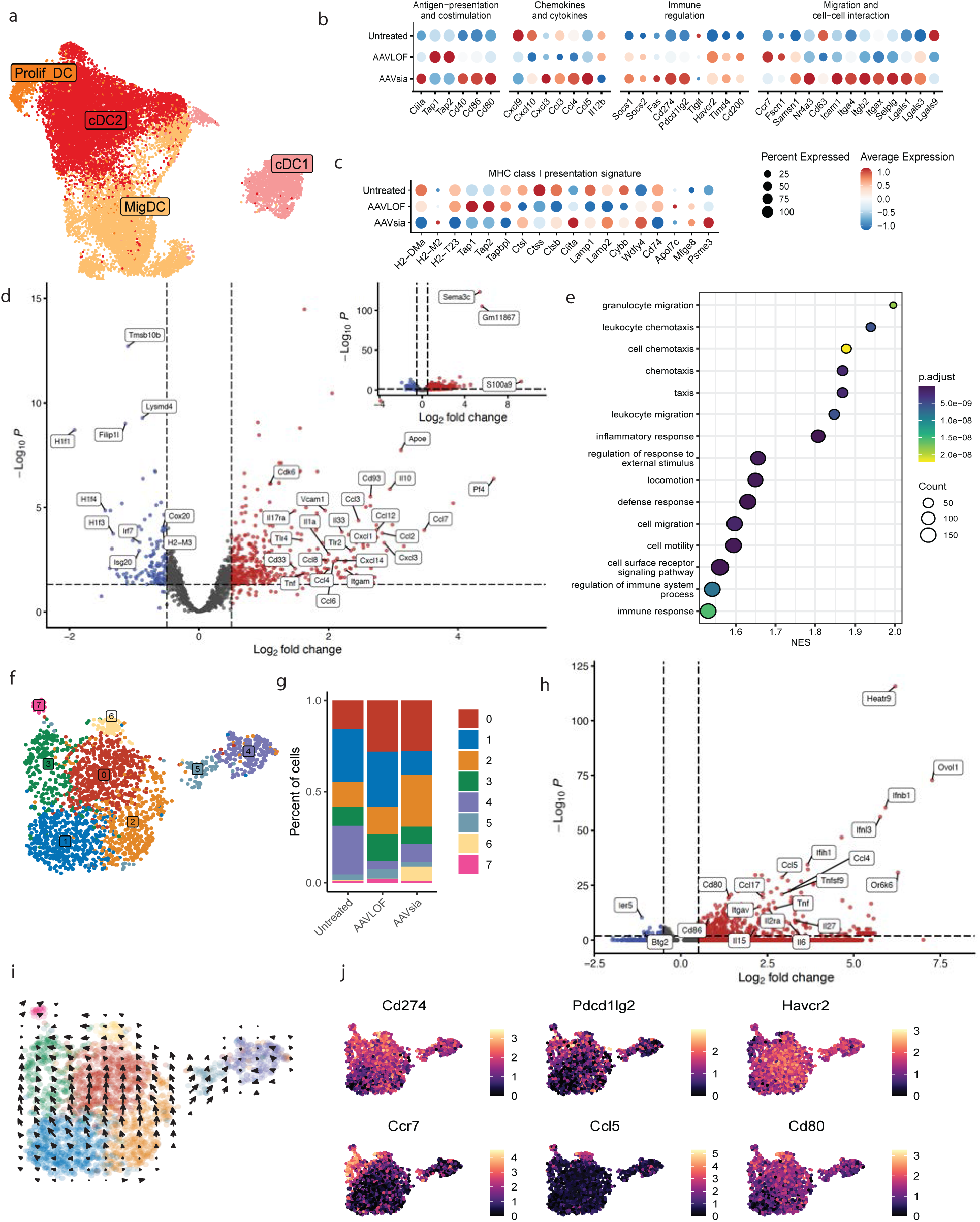
– Desialylation enhances cDC1 activation and promotes a migratory cytokine profile. **a**, Uniform manifold approximation and projection (UMAP) visualization of dendritic cell subsets. **b,c**, Dot-plot showing the expression of selected gene sets in cDC1s, related to antigen presentation and costimulation, chemokine and cytokine signaling, immune regulation, migration and cell–cell interaction, and MHC class I pathways across treatment groups. **d**, Volcano plot depicting pseudo bulked differentially expressed genes (DEGs) in cDC1s following AAV^Sia^ treatment compared with AAV^LOF^. **e**, Pathway enrichment analysis of DEGs in cDC1s from AAVSia- versus AAV^LOF^-treated tumors. **f,g**, Subclustering of the cDC1 population and quantification of subcluster proportions across treatment conditions. **h**, Volcano plot showing DEGs in cDC1 subcluster 6. **i**, RNA velocity analysis embedded on UMAP of the cDC1 population. **j**, Featureplots showing the expression of selected genes across cDC1 subclusters.

Chemokines and migration-related genes were strongly associated with AAV^Sia^ treatment, including *Cxcl3, Ccl3, Ccl4,* and *Ccl5*, as well as genes involved in migration and cell–cell interactions such as *Nr4a3*, a nuclear receptor that regulates surface markers like CCR7 ^24^, which is involved in DC migration. Additionally, *Icam1*, an important adhesion molecule that strengthens the immune synapse with CD8⁺ T cells and is critical for antigen presentation ^25^, was upregulated (**Fig. 4b**). *Cd274*, which encodes the immune checkpoint protein PD-L1, was increased in cDC1 cells upon AAV^Sia^ treatment. PD-L1 is an inhibitory molecule that can limit T cell activation; however, interactions between PD-L1 and CD80 are also important for promoting migration ^26^ (**Fig. 4b**). *Lgals1*, encoding Galectin-1, was increased upon AAV^Sia^ treatment, whereas *Lgals9* (encoding Galectin-9) was decreased. Both galectins often act as negative regulators involved in T cell inhibition ^27, 28^ (**Fig. 4b**).

Further comparison of AAV^Sia^ vs. AAV^LOF^ cDC1 population revealed upregulation of genes related to migration, motility, and inflammatory responses (**Fig. 4d**), which was further confirmed by pathway enrichment analysis (**Fig. 4e**). *Sema3c* was the most differentially expressed gene (DEG) upon AAV^Sia^ treatment and is required for DC migration, particularly during entry into lymphatics ^29^ (**Fig. 4d**).

*Tnf* (TNF-α) is a crucial cytokine for dendritic cell maturation and is essential for priming T cells ^30^, and it was upregulated upon AAV^Sia^ treatment compared with AAV^LOF^. On the other hand, *Il10* coding for IL-10, an anti-inflammatory cytokine, was also upregulated (**Fig. 4d**). However, increasing evidence suggests that IL-10 can induce antitumor effects in an immune-dependent manner. Recent studies have shown that IL-10 receptor signalling on CD8⁺ T cells is required for their activation and proliferation in mouse tumor models ^31^.

cDC1 Subclustering revealed differences in composition among treatment conditions, identifying seven distinct clusters (**Fig. 4f,g and Supplementary Fig. 5e**). Cluster distributions across treatments showed that cluster 6 predominantly appeared upon AAV^Sia^ treatment, whereas cluster 1 was reduced compared to both AAV^LOF^ and AAV^Sia^ conditions (**Fig. 4g and Supplementary Fig. 5e,f**).

Additional comparison of cluster 6 of AAV^Sia^ and AAV^LOF^ treatment groups highlighted differential expression of costimulatory molecules (*Cd80* and *Cd86*), the type I interferon cytokine *Ifnb*, and other inflammatory cytokines such as *Tnf, Il6,* and *Il15* (**Fig. 4h**)*. Tnfsf9* (also known as 4-1BBL) is primarily expressed by activated antigen-presenting cells, including dendritic cells, where it functions as a key costimulatory ligand^32^ (**Fig. 4h**).

RNA velocity analysis of cDC1 subclusters indicated that cluster 1 represents an early population that is reduced in AAV^Sia^-treated mice, whereas cluster 6 corresponds to a more differentiated or mature state (**Fig. 4i).**

UMAP heatmaps confirm that cluster 6 is characterized by co-expression of *Ccr7, Ccl5,* and *Cd80*, while *Cd274* and *Havcr2* are ubiquitously expressed across clusters. *Pdcd1lg2* was more highly expressed in the late-stage cluster 3 (**Fig. 4j**). Importantly, CCR7, CCL5, and CD80 are key molecular makers and regulators involved in dendritic cell maturation, migration, and T cell–stimulatory function, particularly in the context of anti-tumor immune responses^33, 34^.

Similarly to the cDC1 population, migDC also presents increased costimulatory molecules, chemokines, and mainly migration and cell-cell interaction genes (**Supplementary Fig. 6a,b**). The composition of the migDC subclusters differs between the treatment groups, and the RNA velocity analysis shows that cluster 1, corresponding to an early population, was diminished in the AAV^Sia^-treated group, whereas a more mature population (cluster 2) was enriched (**Supplementary Fig. 6c,d,e**).

Taken together, scRNA analysis shows an increased activation, along changes in migration and chemotaxis in cross-presenting DC populations upon AAV^Sia^-treatment.

### Combination glycan engineering shows synergistic effects on cancer progression

In cancer treatment, combination therapies targeting different axes and immune cells often yield higher success rates in overcoming resistance to therapy ^35^. A combination treatment strategy involving checkpoint blockade and fucose targeting together with sialidase treatment, was tested.

Mice were treated with four intra-tumoral injections of AAV^Sia^ combined with anti-PD-1 therapy (**Fig. 5a**). AAV^Sia^ treatment improved the efficacy of anti-PD-1 therapy in MC38 and B16D5 tumor models (**Fig. 5b,c**). EMT6 orthotopic tumors, which are less responsive to anti-PD-1 monotherapy, showed substantial growth inhibition when treated with sialidase alone, and the addition of a PD-1–blocking antibody did not further enhance the therapeutic effect. (**Fig. 5d**).

**Figure 5.**
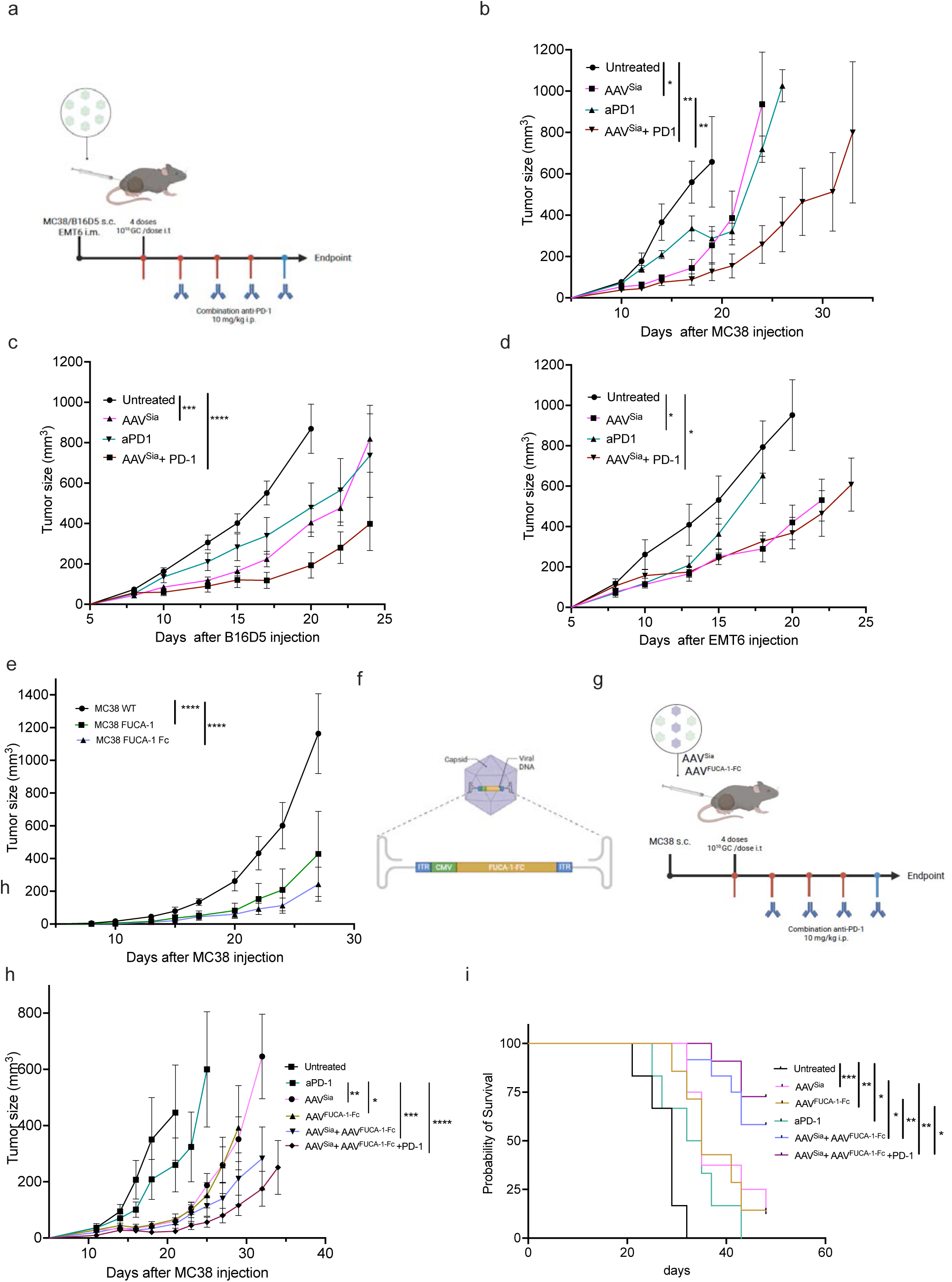
– Combination glycan engineering shows synergistic effects on cancer progression. **a**, Schematic of the treatment strategy combining AAVSia delivery with anti-PD-1 (a-PD-1) immune checkpoint blockade.**b-d**, Tumor growth curves of mice bearing different tumor models treated with AAVSia, a-PD-1, or left untreated (**b**, MC38 n=5-14, **c**, B16D5 n=9-14, and **d**, EMT6 n=8-14) **e**, Tumor growth curves of mice bearing MC38 tumors comparing wild-type ^4^, MC38 cells expressing FUCA-1 (α-L-fucosidase 1), or FUCA-1-Fc fusion protein (n=5-6).**f**, Schematic representation of the engineered AAV vector encoding FUCA-1-Fc (AAV^FUCA-1-Fc^).**g**, Treatment scheme for triple combination therapy using AAV^Sia^, AAV^FUCA-1-Fc^, and a-PD-1 antibody.**h**, Tumor growth curves comparing triple combination therapy with single-treatment controls (n=6-12).**i**, Survival curves for mice receiving triple combination therapy versus single-treatment controls (n=6-12).Growth curves are shown as mean ± S.E.M.Statistical significance among multiple groups was assessed using two-way ANOVA (growth curves), and Tukey’s multiple comparison test. Survival curves were analyzed using the Kaplan–Meier method. *P < 0.05, **P < 0.01, ***P < 0.001. ****P < 0.0001. **a,f,g** were created with BioRender.com.

Analysis of The Cancer Genome Atlas (TCGA) data has revealed elevated expression of fucosyltransferases and sialyltransferases in many cancer types, also correlating with poor survival (**Supplementary Fig. 7a,b**). When both glycosyltransferases are increased in solid cancers, survival rates are significantly decreased (**Supplementary Fig. 7b**). This provided the rationale to combine sialic acid- and fucose-targeting treatment approaches. Small molecule inhibitors, based on substrates of the fucose biosynthetic pathway can be used to inhibit fucosylation ^36^. Recently, a molecule called Fucotrim-1 demonstrated efficacy *in vitro* in prostate cancer cell lines ^37^. To assess the *in vivo* antitumor potential of fucosylation inhibitors Fucotrim-1, an inhibitor of the *de novo* fucose biosynthesis, and B2FF1P^36, 37, 38^, which inhibits fucosylation through the fucose recycling pathway, were investigated. Mice bearing either subcutaneous MC38 colorectal or B16D5 melanoma tumors were intratumorally injected with 10mg/kg Fucotrim-1 or B2FF1P.

Both treatments led to significant tumor growth reduction in two tumor models compared to untreated controls (**Supplementary Fig. 7c,e**). Fucotrim-1 conferred improved survival in both models, but B2FF1P only improved survival in the MC38 model. (**Supplementary Fig. 7d,f**).

We then fused the human fucosidase FUCA-1 (α-L-fucosidase 1) to a human Fc domain (FUCA-1-Fc), containing an antibody-derived signal peptide to enable secretion. The constructs were stably expressed in MC38 cells, which were subsequently injected subcutaneously. MC38 cells expressing FUCA-1 or FUCA-1–Fc exhibited reduced tumor growth (**Fig. 5e**), with the FUCA-1–Fc variant conferring a greater survival benefit (**Supplementary Fig. 7g**). We further engineered an AAV vector encoding FUCA-1 fused to a human Fc domain (AAV^FUCA-1–Fc^; Fig. 5f). Intratumoral administration of AAV^Sia^ and AAV^FUCA-1–Fc^ was combined with intraperitoneal anti–PD-1 checkpoint blockade (**Fig. 5g**). This combination revealed an improved effect between sialidase and fucosidase activity, resulting in increased tumor regression and a higher proportion of tumor-free mice compared to single-agent AAV^Sia^ or AAV^FUCA-1–Fc^ treatments (**Fig. 5h,i**). The addition of anti–PD-1 further enhanced survival, with the triple combination leading to complete tumor clearance in a subset of mice (**Fig. 5i**). Tumor-free mice remained resistant to tumor re-challenge in the contralateral flank, unlike naïve controls, demonstrating the induction of durable antitumor immune memory following local glycan engineering in combination with PD-1 blockade.

These findings suggest that local glyco-engineering could be a new therapeutic approach to enhance classical immune-checkpoint blockade with PD-1-targeting antibodies.

## Discussion

DCs play a pivotal role in priming antitumor T cells by cross-presenting tumor-associated antigens to naïve T cells ^39^. However, the immunosuppressive TME can significantly impact the effector functions of DCs, altering their phenotype and inducing dysfunction and tolerance. Sialylated antigens can induce DC-mediated tolerance by promoting regulatory T cell generation through Siglec-E/9 and by impairing the activity of antigen-specific CD4^+^ and CD8^+^ T cells ^5, 40^. Sialic acids present on DCs themselves can diminish DC-mediated T cell proliferation and activation ^41^ ^42^. Interestingly, the lack of sialic acid, using a sialic acid–blocking mimetic (Ac_5_3F_ax_Neu5Ac), enhanced BMDC and MoDC activation ^43, 44^ and increased antigen-specific CD8⁺ T cell proliferation ^44^. Consistent with this, *in vivo* sialic acid blockade synergized with adoptive transfer of tumor-specific CD8⁺ T cells, promoting DC activation and suppressing tumor growth^45^.

Our study provides evidence that AAV^Sia^-mediated cleavage of sialic acid on the surface of both cancer and immune cells can reduce inhibitory signals in both dendritic cells and T cells, promoting enhanced cross-presentation and subsequent T cell cytotoxicity and tumor control. We have observed a major effect on DCs, however, the immune compartment as a whole could contribute to the maintenance of the anti-tumoral effect. We have shown previously that desialylation also affects macrophage polarization ^6^ and myeloid-derived suppressor cells ^20^. Sialic acid not only binds to Siglec receptors. Indeed, recent findings are demonstrating that other receptors, such as CD28 on T cells, bind to sialylated glycans ^19^, and interestingly, DC-SIGN and DICR bind to nonsialylated glycans on DCs, regulating their function ^46, 47^, suggesting that the use of sialidase as treatment may affect the binding and stability of several receptors.

Systemic administration of sialidase could elicit severe side effects due to its circulation in the bloodstream. Under normal physiological conditions, red blood cells undergo progressive loss of sialic acid, facilitating their clearance from circulation ^48^. Furthermore, desialylation can lead to the clearance of platelets by CD8^+^ T cells in the liver ^49^. A recent study also suggested that the loss of terminal sialylation in glycoproteins present on circulating tumor cell clusters confers an advantage in evading chemotherapy and promotes enhanced metastatic seeding in breast cancer ^50^. This could mean that systemic sialidase treatment could enhance metastatic spread and cancer progression.

To mitigate potential side effects associated with systemic sialidase treatment and reduce the chance of metastasis enhancement, we used a local application. We demonstrated that this intratumoral administration resulted in undetectable levels of sialidase in the bloodstream, reducing potential side-effects and not risking any support of metastatic cancer progression.

Polysialylation, a rare posttranslational modification consisting of long α2,8-linked linear homopolymers of sialic acid, has been shown to control DC trafficking through chemokine CCL21 recognition ^51^. However, its role in directing DC migration to draining lymph nodes during cancer progression remains unclear. In our studies, we observed an increase in migratory DCs within the draining lymph nodes, suggesting that migration was not impaired but rather enhanced, despite influenza sialidase’s broad capacity to cleave all sialic acid linkages.

Targeted sialidases have been used to treat cancer, such as HER2-targeted sialidases and αPD-1-sialidase conjugate, which have shown positive outcomes in mouse models^6,8^. αPD-1-sialidase conjugate has been demonstrated to improve antigen-specific T cell effector function, including in a lymphocytic choriomeningitis virus (LCMV) infection model, and to enhance survival in a murine melanoma model^52^. Another recent study has shown that a bacterial sialidase conjugate (10-1074–SiaD) directs sialidase to the surface of HIV-infected cells via antibody targeting, thereby enhancing immune cell-mediated clearance of these cells ^53^. These findings suggest that sialidase-based approaches may have applications beyond cancer therapy.

Combinatorial treatments are highly beneficial in cancer treatment, using approaches that utilize by different mechanisms, thereby decreasing the likelihood that resistant cancer cells will develop and addressing intrapatient and intratumoral heterogeneity ^9, 54^. Fucosylation is a recognized hallmark of cancer pathogenesis contributing to cancer progression, such as increased cell survival and proliferation, tissue invasion and metastasis ^55^.Our findings demonstrated that the combinatorial approach, cleaving both fucose and sialic acid, was able to improve survival. Combinatorial approaches with classical immune checkpoint inhibitors, such as PD-1 blockade, further enhanced tumor control in mouse models.

A limitation of our study is the lack of human tumor data, highlighting the need for additional studies to investigate expression patterns and potential side effects. The cost of therapeutic AAV vectors limited our ability to test multiple AAV serotypes, and further investigation into more specific promoters is warranted. For example, the AAV vector could be refined using designed ankyrin repeat proteins (DARPins) for targeted expression ^56^. Although our data suggest that dendritic cell desialylation is important for cancer control, the relative contributions of sialic acid loss in different cell types, including tumor cells, remain unclear.

Overall, glycoengineering of the TME was feasible and proved to be beneficial against cancer in preclinical mouse models. This treatment approach could potentially be developed further, and different molecules and/or glycosidases could be expressed to overcome treatment resistance during cancer treatment.

## Materials and Methods

### Cell lines

MC38, MC38-OVA, B16D5 and EMT6 cancer cell lines were kindly provided by Zippelius Laboratory and HEK293T by the Bentires Laboratory, both from the Department of Biomedicine, Basel. Modified MC38 cells were generated by lentiviral transduction as described below. MC38-OVA *B2m*^-/-^ (Knockout) cells were generated using CRISPR/Cas9. The following Guide RNA sequence was used: forward: 5′-TCACGCCACCCACCGGAGAA-3′ and reverse: 5′-TTCTCCGGTGGGTGGCGTGAC-3′, synthesized by Microsynth AG. Lipofectamine CRISPRMAX Cas9 Transfection reagent was used (Invitrogen), according manufacturer’s instructions. After transfection, cell sorting was performed with anti- H-2K^b^ antibody (Biolegend, clone AF6-88.5).

### Lentiviral Contructs

All lentiviral vectors were based on the pLV second-generation lentiviral backbone from VectorBuilder. Proteins were expressed under the control of the truncated EF1A promoter (EFS). The sequence of the H1N1 viral sialidase was based on Uniprot entry C3W5S3. For the FUCA-1 (Uniprot P04066) constructs, the native signal peptide was replaced with a murine IgG heavy chain signal peptide (Uniprot A0N1R4). The FUCA-1-Fc construct additionally contained the human IgG1 CH2/CH3 domains (Uniprot P0DOX5, amino acids 218-449, C222P) at the C-terminus. Lentiviral production and transduction were performed as described previously ^57^.

### Cell culture

Cell lines and primary cells were cultured at 37°C and 5% CO_2_ and regularly checked for mycoplasma contamination. All cell lines except HEKT293T were maintained in Dulbecco’s Modified Eagle Medium (DMEM, Sigma-Aldrich), supplemented with 10% heat-inactivated fetal bovine serum (FBS, PAA Laboratories), 1x MEM non-essential amino acid solution (Sigma) and 1% penicillin/streptomycin (Sigma).

HEK293T, all primary cells and co-cultures were maintained in Roswell Park Memorial Institute Medium (RPMI, Sigma) supplemented with 10% heat-inactivated fetal bovine serum (FBS, Sigma-Aldrich), 1x MEM non-essential amino acid solution (Sigma-Aldrich), 1 mM sodium pyruvate (Sigma), and 1% penicillin/streptomycin (Sigma-Aldrich).

### Mouse strains

C57BL/6N, Balb/c, C57BL/6-Tg(TcraTcrb)1100Mjb/J (OT-I transgenic mice), CD11c-DTR(B6.FVB-17000016L21Rik<Tg(Itgax-DTR7EGFP57Lan>/J) and Batf3^-/-^ (B6.129S(C)-Batf3<TM1KMM>/J) were bred in-house at the University Hospital of Basel (Switzerland). Mice were provided with standard food and water without restriction (License: 1007-2H).

Experiments were performed in accordance with the Swiss federal regulations and approved by the local ethics committee, Basel-Stadt, Switzerland (Approval 3036 and 3099). All animals were bred in-house at the Department of Biomedicine facility (University of Basel, Switzerland) in pathogen-free, ventilated HEPA-filtered cages under stable housing conditions of 45-65% humidity, a temperature of 21-25°C, and a gradual light–dark cycle with light from 7 am to 5 pm.

### Tumor models

WT or transgenic mice were injected subcutaneously into the right flank with 5 × 10⁵ MC38, MC38-OVA, or B16D5 cells in phenol red-free DMEM without additives. EMT6 cell line was injected into the mammary fat pad (5 × 10⁵ cells). Mice were between 8-12 weeks old at the beginning of the experiment.

Tumor size was measured 3 times a week using a caliper. Animals were sacrificed before reaching a tumor volume of 1500 mm^3^ or when they reached an exclusion criterion. Tumor volume was calculated according to the following formula: Tumor volume (mm^3^) = (d^2^*D)/2 with D and d being the longest and shortest tumor parameters in millimeters (mm), respectively.

### *In vivo* Treatments

AAV^Sia^ treatment was initiated when tumors reached a volume of **∼**50 mm³. The virus was diluted in PBS and administered at a concentration of 10¹⁰ genomic copies (GC) per injection, either intravenously (i.v.) or intratumorally (i.t.). Depending on the experiment, 1, 2, or 4 doses were given every 2–3 days. As controls, AAV^LOF^ and/or AAV^null^ were administered under the same conditions.

PD-1 blockade was performed using an anti-PD-1 antibody (clone RMP1-14, BioXCell), administered intraperitoneally (i.p.) in combination with AAV^Sia^ treatment. A total of 4 doses (10 mg/kg each), starting with the second dose of virus, to allow sialidase expression concomitant with the anti-PD-1 expression.

Fucotrim-1 and B2FF1P were synthesized and provided by Dr. Thomas Boltje’s group (Radboud University) ^36, 37, 38^. The compound was diluted in PBS + 15% DMSO and administered i.t. at a concentration of 10 mg/kg when tumors reached 35–75 mm³. Four doses were given every 3–4 days.

FUCA-1 (α-L-fucosidase, gene ID: 2517) was custom-synthesized by GeneUniversal. The enzyme was diluted in PBS and administered i.t. at a concentration of 5 µg/injection when tumors reached 35–75 mm³. Four doses were given every 3-4 days.

### *In vivo* Influenza vaccination

Mice were vaccinated twice subcutaneously with VaxiGripTetra (2021/2022, Sanofi Pasteur, Lot.: V3H691V), containing 15 µg of H1N1 (strain California/2009), before tumor inoculation. The first injection of 1.25 µg was administered 2 weeks before tumor inoculation, followed by a booster of 1.25 µg one week later, resulting in a total of 2.5 µg of inactive H1N1 virus per mouse. The vaccine also contained other influenza strains (H3N2, B Victoria, and B Yamagata). Blood was collected and serum frozen to confirm circulating immunity against H1N1 neuraminidase/sialidase. Tumors were then inoculated, and AAV treatment was performed as described above.

### *In vivo* tumor rechallenge

Long-term tumor-free survival mice were implanted with the same tumor entity in the contralateral flank, with 500,000 MC38 at least 14 days after the primary tumor rejection. Naive C57BL/6 mice were inoculated with MC38 cells, respectively, alongside the rechallenged mice, and tumor growth was monitored until the humane endpoint (<1500 mm^3^).

### Abscopal Tumor Model

Mice were injected subcutaneously with tumors on both flanks, with the right flank serving as the treated tumor site and the left flank inoculated 3–4 days later as the untreated tumor to assess potential abscopal effects. Animals were sacrificed before any tumor reached 1500 mm³ or upon meeting exclusion criteria.

### Tumor digest, splenocyte, serum and PBMC isolation

To obtain single cell suspension, human and mouse tumors were mechanically dissociated and subsequently enzymatically digested using accutase (PAA Laboratories), collagenase IV (Worthington), hyaluronidase (Sigma-Aldrich) and DNase type IV (Sigma) for 1 h at 37°C under constant agitation. Afterwards, samples were filtered using a 100 µM cell strainer and washed. Precision counting beads (BioLegend) were added to all mouse tumors to calculate the number of cells per gram of tumor. Single cells suspensions were used immediately or frozen for later analysis in liquid nitrogen (in 90% FBS and 10% DMSO).

Human peripheral blood mononuclear cells (PBMCs) were isolated from buffy coats by density gradient centrifugation using Hisopaque-1077 (Millipore) and SepMate PBMC isolation tubes (StemCell) according to the manufacturer’s protocol followed by red blood cell lysis using RBC lysis buffer (eBioscience) for 2 min at RT. Subsequently, cells were washed with PBS and were ready co-culture experiments.

For splenocyte isolation, freshly harvested murine spleens were mechanically dissociated by filtering them through a 100 µM filter. After washing, red blood cells were lysed with RBC lysis buffer (eBioscience).

For murine PBMC analysis, blood from the tail vein of mice was collected by tail vein puncture. After washing, red blood cells were lysed as described above, and samples were used immediately for staining.

### Multiparameter flow cytometry

Multicolor flow cytometry was performed on a single cell suspension of cell lines, PBMCs, co-culture system or tumor digest. To avoid unspecific antibody binding, cells were blocked using rat anti-mouse FcγIII/II receptor (CD16/CD32) blocking antibodies (BD Bioscience) for murine and Fc Receptor Binding Inhibitor Polyclonal Antibody (Invitrogen) for human samples, and subsequently stained with live/dead cell exclusion dye (Zombie Dyes, BioLegend). Dextramer staining (mouse MC38 neoantigen, ASMTNMELM) was performed according to the manufacturer’s instructions (Immudex, JA03803) before surface staining.

Surface staining was performed with fluorophore-conjugated antibodies (see Supplementary Tables 1 and 2) or lectins for 30 minutes at 4°C in FACS buffer (PBS, 2% FCS, 0.5 mM EDTA).

Stained samples were fixed using IC fixation buffer (eBioscience) until further analysis. For intracellular staining, cells were fixed and permeabilized using the Foxp3/transcription factor staining buffer set (eBioscience) and 1x Permeabilization buffer (eBioscience) according to the manufacturer’s instruction (see Supplementary Tables 1 and 2). Compensation was performed using AbC Total Antibody Compensation Bead Kit (Invitrogen) or cells.

Samples were acquired on LSR II Fortessa flow cytometer (BD Biosciences) or Cytek Aurora (Cytek Biosciences) and analyzed using FlowJo 10.10 (BD Bioscience). Cell sorting was performed using a BD FACSAria III or BD FACSMelody (BD Bioscience). Doublets, cell debris and dead cells were excluded before performing downstream analysis. Fluorescence-minus-one (FMO) samples were used to define the gating strategy and calculate mean fluorescence intensity (MFI).

### OT-I cells

OT-I CD8^+^ T cells were isolated from the spleen of 7-10 week-old female C57BL/6-Tg(TcraTcrb)1100Mjb/J (OT-I) mice using the EasySep CD8^+^ T-cell isolation kits (STEMCELL), according to the manufacturer’s instructions, leading to a CD8^+^ T-cell purity of around 96%. T cells were co-cultured in RPMI 1640 medium supplemented with 10% FBS, L-glutamine, non-essential amino acids (0.1 mM, Sigma) and sodium pyruvate (1 mM, Gibco) and IL-2 (200IU/ml, Iovance).

### Recombinant adeno-associated viruses

Neuraminidase 1 derived from *Influenza A* (AAV^Sia^, Gene ID: 23308118), a loss-of-function mutant of the same enzyme (AAV^LOF^, Y402F), or human α-L-fucosidase (AAV^FUCA^, Gene ID: 2517) was each cloned into an AAV serotype 2 (AAV2) vector. All recombinant virus cloning and production were performed by Vector Biolabs. AAV^null^ control vectors (Vector Biolabs #7026), containing a CMV promoter but no transgene, were used as ready-to-use empty vector controls. In short, HEK293 were co-transfected with AAV plasmid with the Gene-of interest (GOI) under the CMV promoter, together with the replication helper-plasmid DNA comprising AAV2 capsid and replication genes. In the AAV plasmid, the REP and CAP genes of wild type AAV were deleted, so there are only 2 copies of ITRs (∼145 bp/each) left. 2 days after transfections, cell pellets are harvested, and viruses are released through 3x cycles of freeze/thaw. Viruses are purified through CsCl-gradient ultra-centrifugation, followed by desalting. Viral titer (GC/ml - genome copies/ml) is determined through real-time PCR. The purity of AAV proteins is analyzed using SDS-gel silver staining. Only AAV preps with >90% purity can pass the QC.

### Monocyte-derived dendritic cells (moDC) generation and co-culture

For monocyte isolation, CD14-positive cells were magnetically isolated using a human CD14-positive selection kit (StemCell) according to the manufacturer’s instructions. The isolated cells were resuspended in fresh complete RPMI medium and incubated for 7 days with GM-CSF (20ng/ml, Peprotech) and IL-4 (10ng/ml, Peprotech) for moDcs generation. LPS 100 ng/ml (Sigma-Aldrich) was used as a maturation stimulus for 24h. After the incubation period, moDCs were either pre-treated for 20 minutes with recombinant active *influenza A H1N1* neuraminidase/sialidase (10U/ml, Sino Biological) or left untreated, and then co-incubated with T cells that had been magnetically isolated using a negative selection kit (StemCell) from a different healthy donor, also either pre-treated with sialidase or left untreated. moDCs and T cells were cultured for 5 days (ratio 1:2), after which flow cytometry was performed to analyze T cell proliferation and activation.

### Cytokine and chemokine analysis

Murine serum samples from various treatment groups were thawed on ice, and aliquots were shipped on dry ice to Eve Technologies (Canada) for analysis. Cytokine and chemokine concentrations were measured and calculated by Eve Technologies. For data visualization, normalized values (z-scores) were calculated for each cytokine based on the mean and standard deviation of that marker.

### Immunofluorescence

For the analysis of sialic acid binding or expression of H1N1 sialidase by immunofluorescence, frozen sections of tumors embedded in optimal cutting temperature (OCT) compound were cut and prepared using a cryostat. Samples were stored at –80 °C until staining, which was performed immediately after fixation with ice-cold methanol for 5 minutes. To detect sialic acid, SNA-Cy3 (Vector Laboratories) was used at 10 µg/ml and incubated in PBS for 2h at room temperature. To detect H1N1 neuraminidase/sialidase, the fixed samples were blocked with 1% BSA for 1 h, followed by incubation with a polyclonal anti-neuraminidase/sialidase antibody (LS-C486838, LS-Bio) as the primary antibody (1:500) overnight at 4 °C. After washing three times with PBS-T, the secondary antibody anti-rabbit A647 (A21244, Invitrogen) was added (1:1500) for 2h at room temperature. After incubation, slides were washed and mounted with ProLong Diamond Antifade Mountant containing DAPI. Images were acquired using a Nikon Eclipse Ti-S microscope and analyzed with QuPath.

### ELISA

Mouse serum was used to detect either circulating Influenza A H1N1 sialidase/neuraminidase or anti-H1N1 sialidase/neuraminidase antibodies upon treatment. To detect sialidase in serum, high-binding ELISA microplates (Corning) were coated with 50 μl/well of serum diluted in 50 μl of coating buffer and incubated overnight at 4 °C. Plates were blocked with 1% BSA for 2 h at room temperature. Wells were then washed three times with PBS-T (PBS + 0.05% Tween-20) and incubated with a biotinylated anti-H1N1 neuraminidase antibody (BAF4858, R&D Systems) diluted 1:2000 in blocking buffer. As a control, a standard curve was generated using serial concentrations of inactive H1N1 sialidase/neuraminidase (11058-V07B, Sino Biological). After washing, Streptavidin-HRP (1:2500, BD Biosciences) was added and incubated for 1h at room temperature. Plates were washed again, and TMB substrate (BD Biosciences) was added and incubated for ∼20 minutes at room temperature. The reaction was stopped with stop solution, and the absorbance was measured immediately at 450 nm. To detect antibodies in the serum of vaccinated animals, serum samples were diluted in coating buffer and added into plates at different concentrations overnight. Washing and blocking were performed as described above. Plates were then incubated with 20 ng/mL of inactive H1N1 sialidase/neuraminidase (Sino Biological) for 2 h at room temperature. After washing, the protocol was continued using biotinylated anti-H1N1 neuraminidase/sialidase, HRP-conjugated secondary antibody, and TMB substrate as described above.

### Cross-presentation assay

Bone marrow cells from C57BL/6 mice were differentiated into dendritic cells (DCs) by culture with Flt3L (150 ng/ml, Peprotech) for 9 days, with half of the medium replaced every 2 days. XCR1⁺ DCs (cDC1s) were enriched using magnetic beads with an XCR1-PE antibody (Clone: ZET) according to the manufacturer’s instructions for the EasySep PE Positive Selection Kit. Purified XCR1⁺ DCs were seeded at a density of 40,000 cells per well in a U-bottom 96-well plate. TNF-α (5 ng/ml, Peprotech) was added to activate DCs.β2m⁻/⁻ MC38-Ova tumor cells were irradiated with 100 Gy at a density of 10,000 cells per well and co-cultured with the purified DCs overnight. CTV-labeled (Thermo Fisher) CD8⁺ T cells from OT-I mice were then added at a density of 100,000 cells per well. After 72 h, T cell proliferation was analyzed by flow cytometry based on CTV dilution and surface marker staining.

Tumor XCR1⁺ cells were harvested from differently treated MC38-OVA tumors using anti-XCR1 microbeads (Miltenyi Biotec), seeded at a density of 40,000 cells per well, and co-cultured with 100,000 CTV-labeled OT-I cells for 72 hours to assess T cell proliferation and cross-presentation.

### Virus quantification by PCR

Host genomic DNA and viral DNA were isolated from snap-frozen treated or untreated tissues using QIAamp DNA Mini Kit (Qiagen) according to the manufacturer’s protocol. DNA yield and quality were analyzed by Nanodrop. The number of AAV GC per ng of host DNA was determined by a standard curve, which was generated using the unmodified AAV2-CMV plasmid DNA (Vector Biolabs) extracted by degradation of the viral capsid at 95°C for 10 minutes, and standards were prepared by 10-fold serial dilutions. Samples obtained from untreated mice were used for minimal threshold detection of the viral DNA. Three independent qPCR reactions were performed on Applied Biosystems 7500 FAST Real-Time PCR System using cytomegalovirus (CMV) promoter as a target sequence. Detection CMV primers (forward: 5’-CCC ACT TGG CAG TAC ATC AA-3’, reverse: 5’-GCC AAG TAG GAA AGT CCC ATA A-3’) and the custom Taqman probe (5’-FAM-CAT AAT GCC AGG CGG GCC ATT TAC-3-BHQ-1’) were synthesized by Microsynth. All of the PCRs were performed in a 20 μL final volume with 900 nM of each primer, and 300nM probe, 100 ng of diluted sample DANN, and 2X GoTaq Probe qPCR Master Mix (Promega). All of the reactions were performed in triplicate, and a negative control (template DNA) was included. FAST cycling qPCR was performed with 1 cycle of 2 minutes at 95 °C to activate DNA polymerase, followed by 45 cycles of 3-second denaturation at 95 °C and 30-second annealing at 60 °C.

### Cell Sorting and cell-hashing for ScRNA-seq

MC38-OVA cells (5 × 10⁵) were inoculated subcutaneously into mice. AAV treatment was initiated when tumors reached a volume of 35–75 mm³. Two doses were administered 3 days apart. One week after the first injection, tumors were harvested and enzymatically digested. Cells were stained with CD11c-APC (BD Biosciences) and magnetically enriched for CD11c using the APC positive selection kit (StemCell).

Cells were then processed for CITE-seq using TotalSeq antibodies against XCR1 and CCR7 (BioLegend). In parallel, samples were stained for cell sorting with F4/80-FITC, MHCII-Pacific Blue, and Zombie NIR live/dead dye. Sorting was performed by excluding F4/80⁺ macrophages and dead cells, and gating on dendritic cells (CD11c⁺MHCII⁺).

### scRNA-seq library preparation

Single-cell 3′ gene expression profiling on the single-cell suspensions using the Chromium Next GEM Single Cell 3’ with Feature Barcode technology from 10x Genomics was performed according to the manufacturer’s instructions. Cells were loaded in 10x Genomics cartridge to target recovery of approximately 20,000 cells for each condition. Mouse specific cell hashing with TotalSeq-B hashtag antibodies (Biolegend) was performed according to supplier instructions. Cell-barcoded 3′ gene expression libraries and feature barcode libraries were sequenced on Illumina NovaSeq6000 system. Resulting per condition reads where demultiplexed per mouse and mapped to the GRCm39 murine reference genome using CellRanger v8.0.1 (10x Genomics).

### scRNA-seq data analysis

Read count matrices were processed and analyzed in R v.4.4.2 using Seurat v.5.2.0 with default parameters in all functions, unless other specified. Cell quality was investigated and low-quality cells were removed based on nFeatures under 1000, nCounts under 1500, mitochondrial gene percentage of more than 6% and ribosomal gene percent of less than 5%. Subsequently, data was normalized variable features identified and scaled. From variable features irrelevant genes like histone, mitochondrial genes or ribosomal gens were excluded using the gene blacklist for mice published at https://github.com/Japrin/scPip/. Mitochondrial gene percentage as well as cell cycle was regressed out during scaling using the murine cell cycle gene set published at https://github.com/Japrin/scPip/. After PCA reduction, the first 15 PCs were used for neighbor calling with a resolution of 0.6 for cluster identification. UMAP was also calculated on the first 15 PCs. Clustering was repeated for the subsets cDCs, cDC1 and MigDC using the same approach. For this also ribosomal percentage and Browne interferon response genes were regressed during scaling and the first 20 PCs were used for clustering at a resolution of 0.8. For data visualization the R packages ggplot2 v3.5.1, Nebulosa v1.14.0, dittoSeq v1.16.0, enrichplot v1.24.4 and EnhancedVolcano v1.22.0 were used.

### Differential Gene Expression Analysis

To compare expression differences between conditions in cDC1s a pseudo bulk approach was used to reflect individual mice as replicates. First, count data was aggregated per mouse. Then, to perform DGE between conditions the standard edgeR v4.2.2 workflow was followed using the Benjamini-Hochberg Procedure for p-value correction. Cluster specific gene expression was identified using the FindAllMarkers or FindMarkers functions implemented in Seurat.

### Gene set enrichment analysis

The in Seurat implemented FindMarkers function was used to receive up and downregulated markers between conditions. For this the logfc.threshold was set to zero. With the R package clusterprofiler v4.12.62 gene set enrichment analysis of the Gene Ontology database was performed. The results were ordered based on normal enrichment score and only highest significant ontologies were used for data interpretation.

### RNA velocity trajectory analysis

Spliced and unspliced reads were counted using the velocyto.py package v0.17.17 from aligned bam files generated by CellRanger. The resulting count matrices where normalized and the top 2000 variable genes with a minimum of 20 shared counts were selected. Moments were calculated for velocity estimation and used in the dynamic model of scVelo to learn the unspliced/spliced phase trajectory. For visualization, the resulting trajectory vectors were embedded onto the UMAP space.

### scRNAseq Data availability

The scRNAseq data generated in this study is available at Gene Expression Omnibus (GEO) accession no. GSE292054.

### Statistics

For statistical considerations, except ScRNAseq, GraphPad Prism was used (Version 10). For efficacy analysis, including progression-free survival and overall survival, Kaplan-Meier curves were used. Specific tests are described in the legends of the figures. Differences were considered statistically significant when p-values were below 0.05.

## Contributions

HL and NRM planned the project and experimental design. NRM, FF, AZ, IMS,JFAP, MS, ACP, and ER performed and analyzed experiments. NRM,TB, and HL interpreted the results. MS processed, analyzed, and interpreted scRNAseq data. HL and NRM wrote the manuscript. All authors reviewed the manuscript.

## Artificial Intelligence Statement

Grammarly and ChatGPT were used to proofread the text for typos, grammar, and syntax errors. No generation of content was performed.

