## Supplemental Data for "Local glycan engineering induces systemic antitumor immune reactions via antigen cross-presentation"

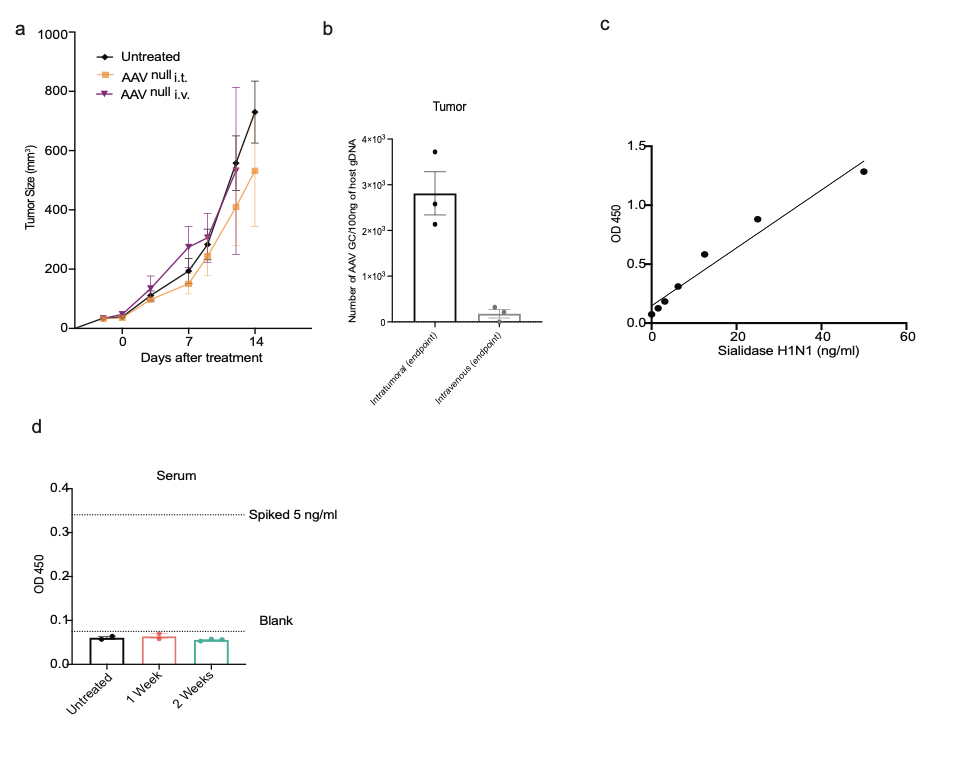

**Supplementary Figure 1**- **a**, MC38 cells were injected subcutaneously (s.c.) into mice and treated with AAV^Sia^ either intratumorally (i.t.) or intravenously (i.v.), left untreated or treated with AAV^null^ . **b,** Quantitative PCR (qPCR) was performed to detect AAV at endpoint in tumors upon intratumoral or intravenous treatment.**c,d**, ELISA was performed to detect Sialidase H1N1 in the serum of treated mice, standard curve, and levels of sialidase detected in the serum upon intratumoral treatment. Growth curves are shown as mean ± S.E.M. Bar plots are presented as mean ± S.D.

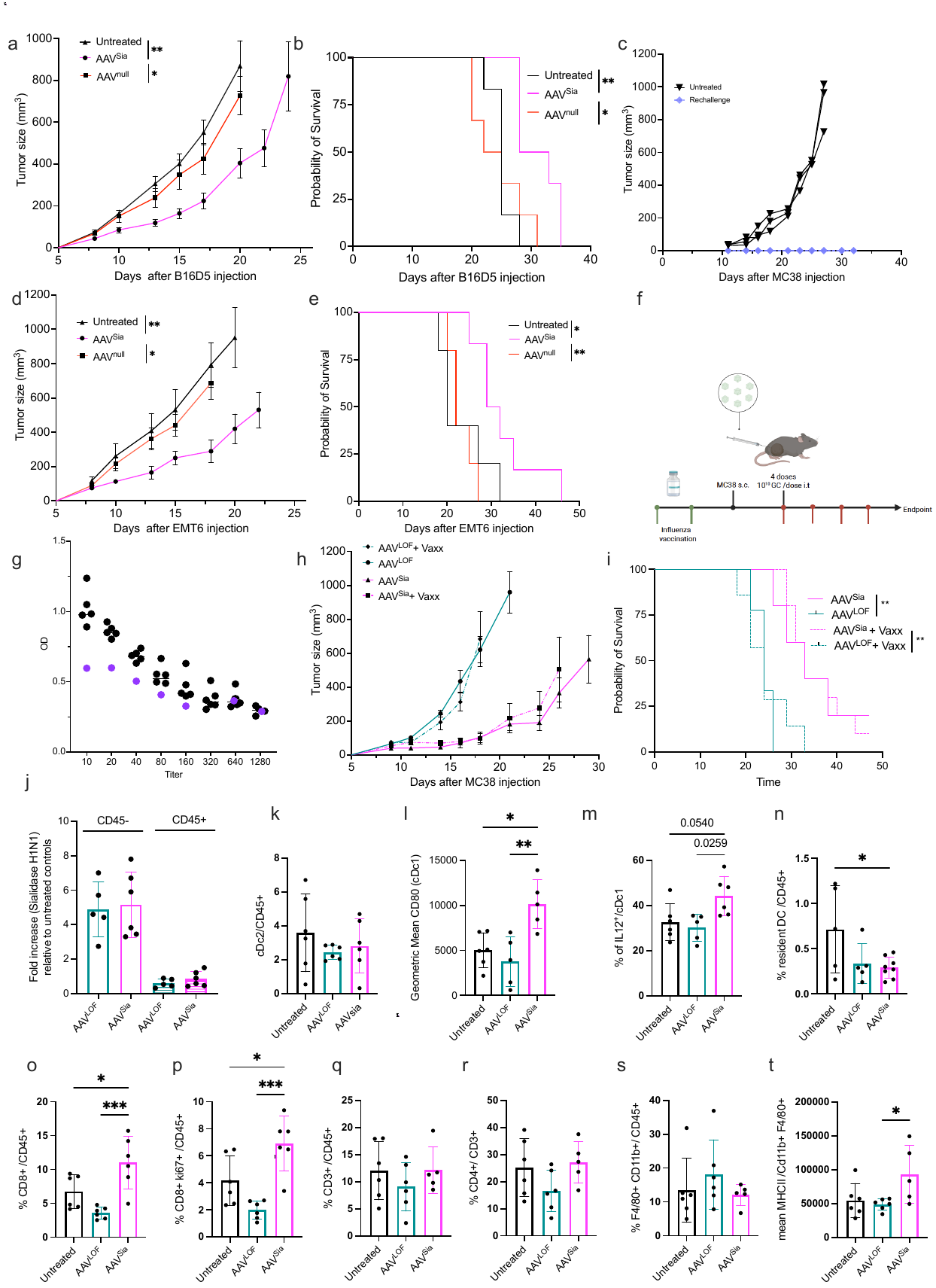

**Supplementary Figure 2-** **a,b**, B16D5 tumor growth and survival curves after four i.t doses of AAV^Sia^ and AAV^null^. **c**, Re-challenge experiment in tumor-free mice following AAV^Sia^ treatment; black curves represent newly injected mice, and purple curves represent previously tumor-free mice. **d,e**, EMT6 tumor growth and survival curves after four i.t doses of AAV^Sia^ and AAV^null^. **f**, Treatment scheme showing influenza vaccination followed by MC38 tumor injection and subsequent AAV^Sia^ or AAV^LOF^ treatment. **g**, Antibody titers in the blood of influenza-vaccinated mice (black dots) compared with a non-vaccinated mouse (lilac dot). **h,i**, Tumor growth and survival curves of vaccinated mice treated with AAVLOF or AAVSia. **j**, H1N1 sialidase expression in CD45+ and CD45− tumor cell populations after short-term treatment (2 injections). **k**, Intratumoral conventional dendritic cell type 2 (cDC2) frequency. **i**, Geometric mean of CD80 expression in the cDC1 population. **m**, Intratumoral frequency of IL-12+ cDC1 cells. **n**, Frequency of resident dendritic cells (mDCs) in the draining lymph node. **o**, Intratumoral CD8+ T cell frequency. **p**, Frequency of CD8+ T cells positive for the proliferation marker Ki67. **q**, CD3+ cell frequency. **r**, CD4+ T cell frequency. **s**, Frequency of F4/80+ CD11b+ macrophages. **t**, Mean MHC II expression in the macrophage population. Growth curves are shown as mean ± S.E.M. Bar plots are presented as mean ± S.D. Statistical significance among multiple groups was assessed using one-way ANOVA (bar plots) or two-way ANOVA (growth curves). Survival curves were analyzed using the Kaplan–Meier method. *P < 0.05, **P < 0.01, ***P < 0.001.

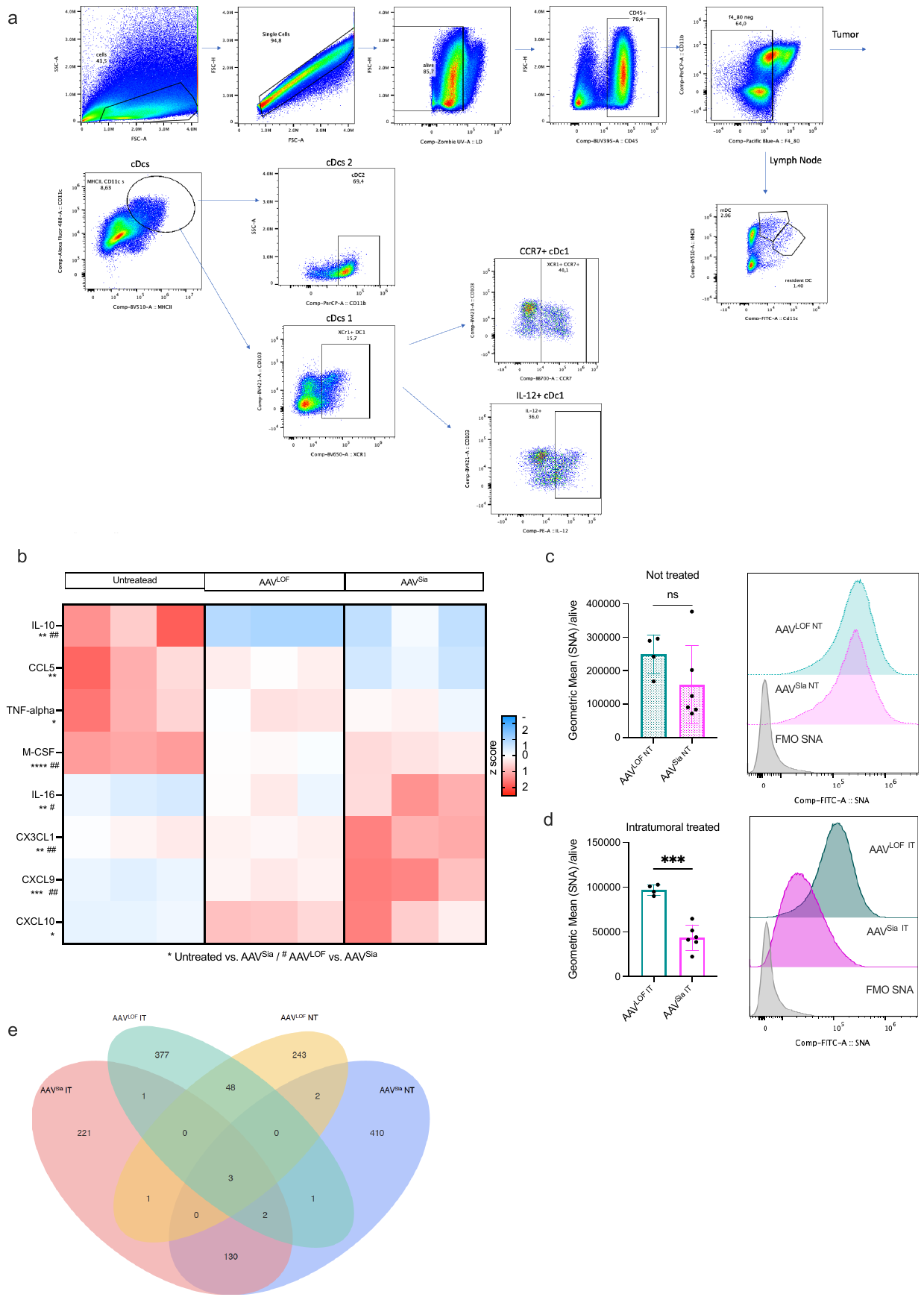

**Supplementary Figure 3** a, Gating strategy for intratumoral and lymph node dendritic cells. b, Heatmap of cytokines and chemokines that are statistically significant following short-term AAV^Sia^ or AAV^LOF^ treatment. c,d, SNA geometric mean fluorescence intensity in live intratumoral cells from contralateral non-treated tumors (NT) or intratumorally treated (IT) tumors with AAV^Sia^ or AAV^LOF^, with representative histogram plots. e, Venn diagram of clonotypes (defined by amino acid sequence) in an abscopal model following intratumoral (IT) treatment with AAV^Sia^ or AAV^LOF^, compared with contralateral non-treated tumors. Statistical significance among multiple groups was assessed using one-way ANOVA . # or*P < 0.05, ## or**P < 0.01, ### or ***P < 0.001.

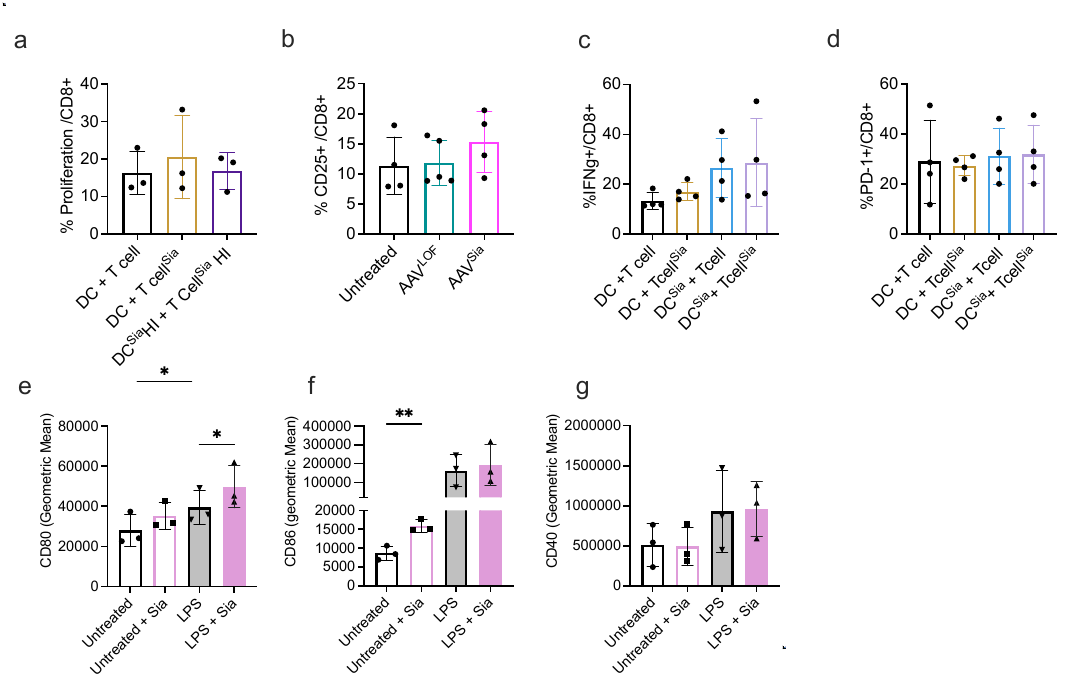

**Supplementary Figure 4- a,** Quantification of T cell proliferation in the in vitro assay following pre-treatment with soluble H1N1 sialidase (Sia) or heat-inactivated sialidase (Sia HI). **b,** Frequency of CD25⁺ cells in the in vitro cross-presentation assay using XCR1⁺ intratumoral cells co-incubated with OT-I CD8⁺ T cells. **c,** Frequency of interferon-γ–producing CD8⁺ T cells in the mixed lymphocyte reaction (MLR) assay. **d,** Frequency of PD-1⁺ CD8⁺ T cells in the MLR assay. **E-g,** Geometric mean fluorescence intensity (gMFI) of CD80 (**e**), CD86 (**f**), and CD40 (**g**) in monocyte-derived dendritic cells (moDCs) following pre-treatment with sialidase and overnight maturation with lipopolysaccharide (LPS). Bar plots represent mean ± s.d. (n = 3-4). Statistical significance among multiple groups was assessed by one-way ANOVA with Tukey’s multiple-comparison test (**a–d**). Unpaired t-tests were used for pairwise comparisons (Untreated vs. Sia + LPS, and LPS vs. Sia + LPS) (**e–g**). *P < 0.05, **P < 0.01, ***P < 0.001.

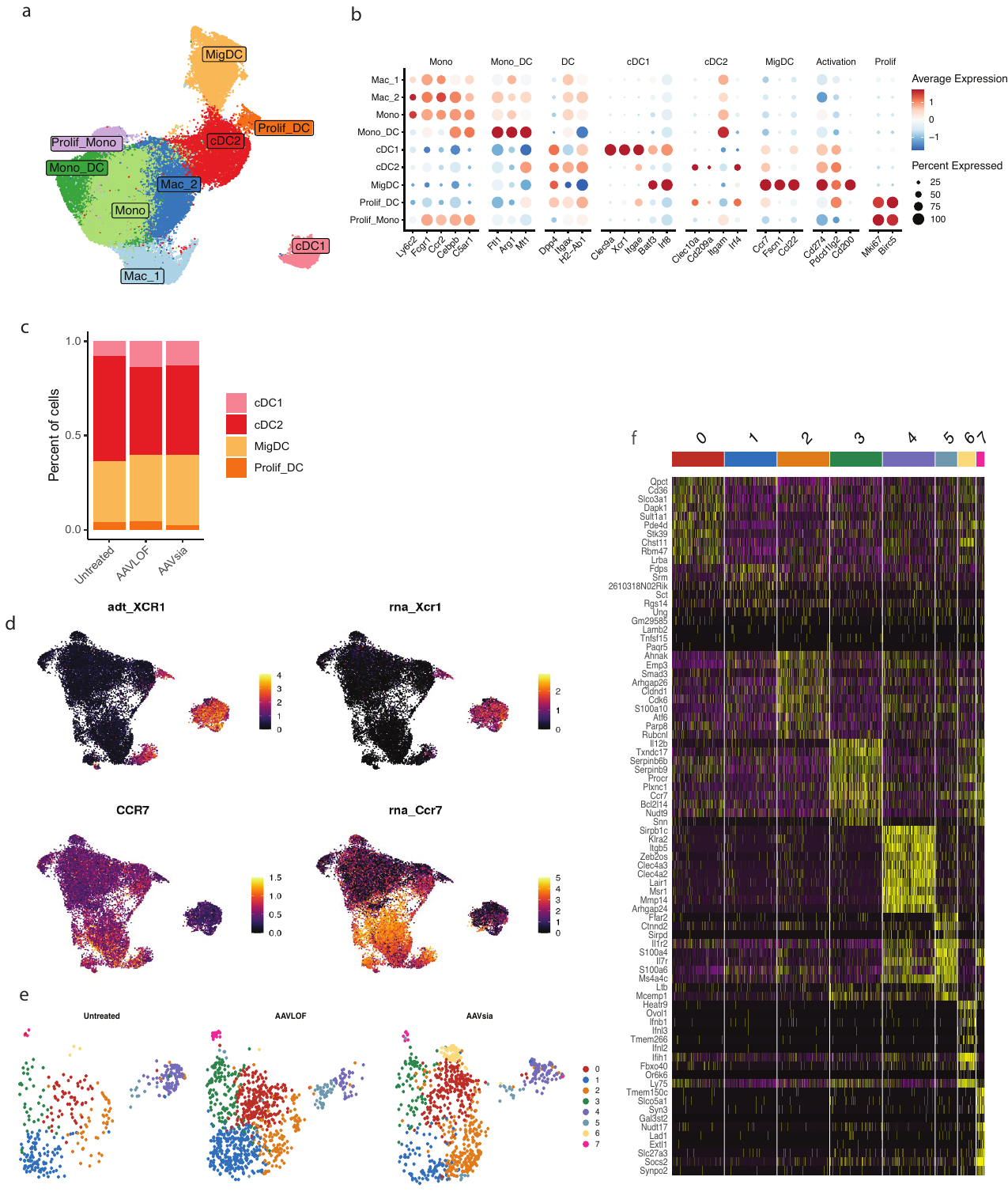

**Supplementary Figure 5-** **a**, Uniform manifold approximation and projection (UMAP) of CD11c⁺ MHCII⁺ F4/80⁻ sorted immune cell subpopulations. **b**, Dotplot showing annotated immune populations and a selection of their associated marker genes. **c**, Proportions of dendritic cell subsets across treatment groups. **d**, Featureplot representation of selected markers showing protein expression of XCR1 and CCR7 and comparison with transcriptomic expression. **e**, UMAP of the cDC1 subset stratified by treatment group. **f**, Heatmap showing top gene expression per cDC1 subclusters.

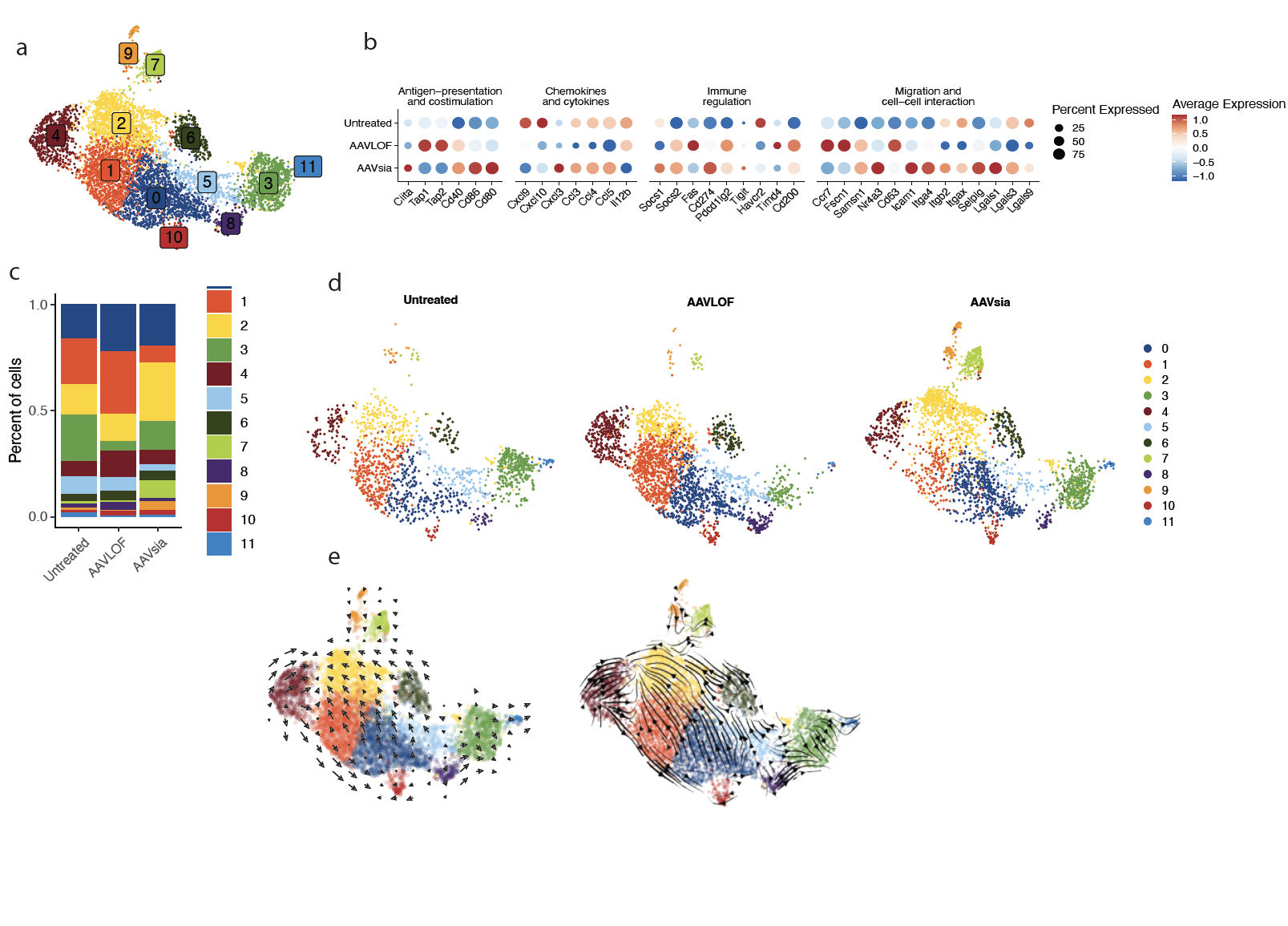

**Supplementary Figure 6-** **a**, Uniform manifold approximation and projection (UMAP) showing subclustering of migratory dendritic cells (migDCs). **b**, Dotplot illustrating the expression of selected gene sets in migDCs, including signatures associated with antigen presentation and costimulation, chemokine and cytokine signaling, immune regulation, migration, and cell–cell interaction. **c**, Proportions of migDC subsets across treatment groups. **d**, UMAP of migDC subsets stratified by treatment group. **e**, RNA velocity analysis of migDC subsets embedded on UMAP space.

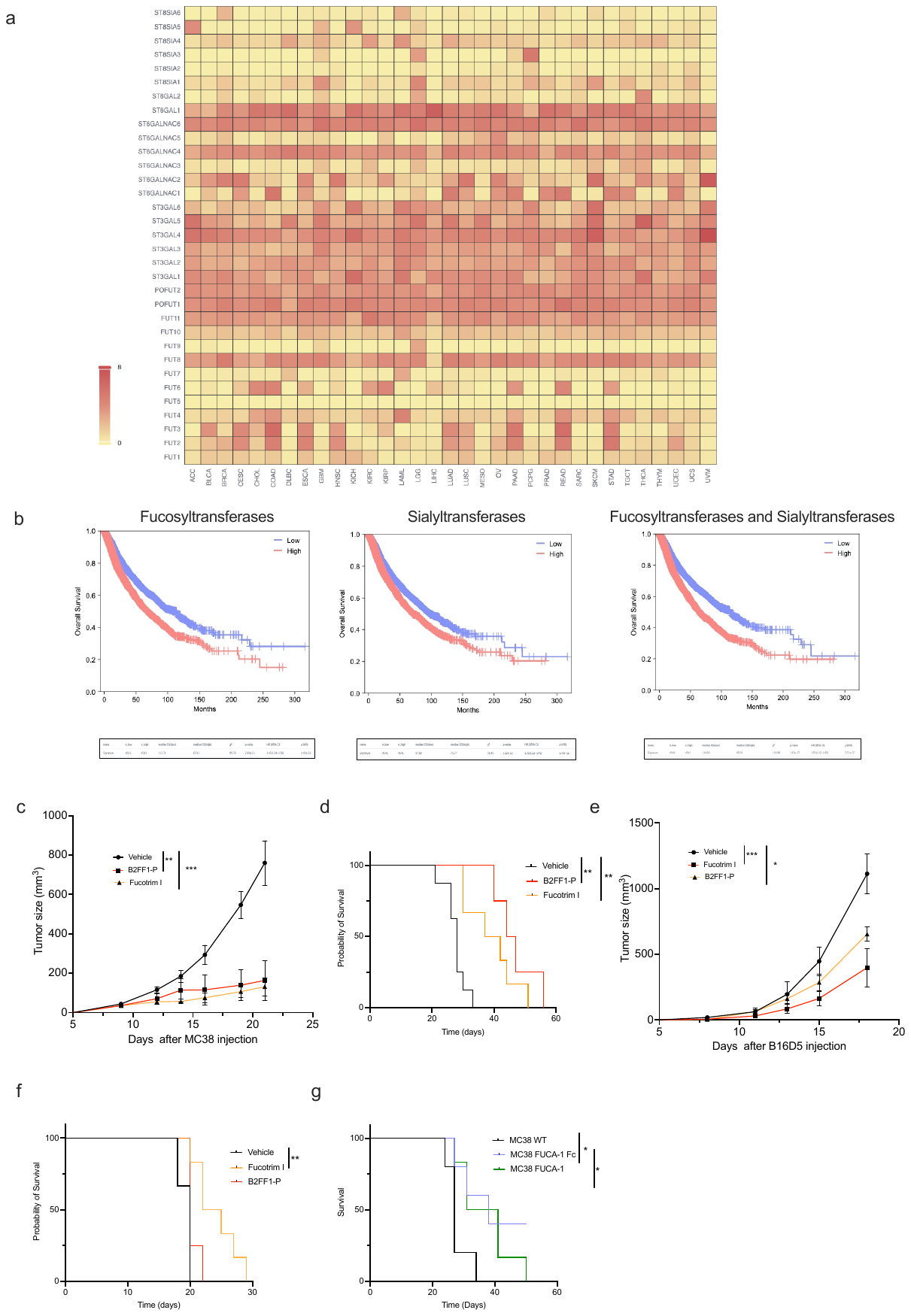

**Supplementary Figure 7-** **a**, Gene expression analysis of sialyltransferases and fucosyltransferases across multiple cancer types using the GEPIA3 web server, based on data from The Cancer Genome Atlas (TCGA).**b**, Overall survival analysis comparing high versus low expression (50/50 stratification) of sialyltransferases, fucosyltransferases, or their combined expression across all solid tumor types in the TCGA dataset, performed using GEPIA3.**c,d**, MC38 tumor-bearing mice treated with four intratumoral injections of Fucotrim-I, B2FF1-P, or vehicle (PBS + 20% DMSO):**c**, tumor growth curves; **d**, survival curves. **e,f**, B16D5 tumor-bearing mice treated with four intratumoral injections of Fucotrim-I, B2FF1-P, or vehicle (PBS + 20% DMSO):**e**, tumor growth curves; **f**, survival curves.**g**, Survival curves of mice bearing MC38 tumors comparing wild-type (WT), FUCA-1-expressing, or FUCA-1-Fc-expressing cells. Growth curves are shown as mean ± S.E.M. Statistical significance among multiple groups was assessed using two-way ANOVA (growth curves) or Kaplan-Meier (survival curves). *P < 0.05, **P < 0.01, ***P < 0.001.

**Supplementary Tables**

|  | **Colour** | **Clone** | **Dilution** | **Supplier** | **Cat. Number** |
| --- | --- | --- | --- | --- | --- |
| **Live-dead** | Zombie UV |  | 100 | Biolegend | 423107 |
| **Live-dead** | NIR |  | 500 | Biolegend | 423105 |
| **CD45** | BUV395 | 30F-11 | 100 | BD Biosciences | 564279 |
| **CD4** | BUV496 | GK1.5 | 100 | BD Biosciences | 612952 |
| **Ly-6G** | BUV563 | 1A8 | 200 | BD Biosciences | 612921 |
| **NKp46** | BUV661 | 29A1.4 | 70 | BD Biosciences | 741678 |
| **CD3** | BUV805 | 145-2C11 | 70 | BD Biosciences | 749276 |
| **CD103** | BV421 | 2E7 | 150 | Biolegend | 121442 |
| **CCR7** | BB700 | 4B12 | 100 | eBioscience | 566462 |
| **CD8** | PE-Cy5 | 53-6.7 | 100 | eBioscience | 480081 |
| **MHCII** | BV510 | M5/114.15.2 | 300 | Biolegend | 107635 |
| **CD80** | BV605 | 16-10A1 | 70 | Biolegend | 104729 |
| **XCR1** | BV650 | ZET | 100 | Biolegend | 148220 |
| **PD-1** | BV785 | 29F.1A12 | 100 | Biolegend | 135225 |
| **CD19** | BV570 | 6D5 | 100 | Biolegend | 115535 |
| **CD11c** | FITC | N418 | 100 | Biolegend | 117306 |
| **Tim-3** | BV711 | RMT3-23 | 100 | Biolegend | 119727 |
| **CD25** | PE-Cy5.5 | PC61.5 | 100 | eBioscience | 35025182 |
| **CD25** | APC | PC61.5 | 200 | Biolegend | 102012 |
| **CD69** | PercP-Cy5.5 | H1.2F3 | 200 | Biolegend | 104522 |
| **F4/80** | Pacific Blue | BM8 | 100 | Biolegend | 123124 |
| **CD11b** | PercP | M1/70 | 100 | Biolegend | 101229 |
| **Ki67** | AF532 | SolA15 | 200 | eBioscience | 58569882 |
| **TCF-7/TCF1** | AF700 | 812145 | 100 | R&D sytems | FAB82224N |
| **GzmB** | PE-eFluor610 | NGZB | 100 | eBioscience | 61889882 |
| **FoxP3** | AF660 | FJK-16s | 100 | eBioscience | 606577382 |
| **IL-12** | PE | C15.6 | 50 | BD Biosciences | 554473 |

Table 1- Dye and anti-mouse conjugated antibodies for flow cytometry

|  | **Colour** | **Clone** | **Dilution** | **Supplier** | **Cat. Number** |
| --- | --- | --- | --- | --- | --- |
| **Live-dead** | Aqua |  | 200 | ThermoFisher | L34957 |
| **CD8** | BUV805 | RPA-T8 | 400 | BD Biosciences | 568334 |
| **IFN-gamma** | BB700 | B27 | 100 | BD Biosciences | 566395 |
| **PD-1** | PE-Cy7 | EH12.1 | 50 | BD Biosciences | 561272 |
| **TCF-1/7** | AF647 | 7F11A10 | 50 | Biolegend | 655204 |
| **GzmB** | AF700 | QA16A02 | 400 | Biolegend | 372222 |
| **CD107a** | APC-H7 | H4A3 | 200 | BD Biosciences | 561343 |
| **CD40** | FITC | 5C3 | 100 | Biolegend | 334306 |
| **CD80** | BV605 | 2D10 | 50 | Biolegend | 305225 |
| **CD86** | BV711 | IT2.2 | 100 | Biolegend | 305440 |
| **CD3** | BUV395 | SK7 | 200 | BD Biosciences | 564001 |
| **CD4** | PE | SK3 | 100 | eBioscience | 12004742 |

Table2- Dye and anti-human conjugated antibodies for flow cytometry
